# Self-regulating arousal via pupil-based biofeedback

**DOI:** 10.1101/2022.08.26.505388

**Authors:** Sarah Nadine Meissner, Marc Bächinger, Sanne Kikkert, Jenny Imhof, Silvia Missura, Manuel Carro Dominguez, Nicole Wenderoth

**Affiliations:** Neural Control of Movement Laboratory, Department of Health Sciences and Technology, ETH Zurich, Technoparkstrasse 1, 8005 Zurich, Switzerland; Neuroscience Center Zurich, University and ETH Zurich, Winterthurerstrasse 190, 8057 Zurich, Switzerland; Future Health Technologies, Singapore-ETH Centre, Campus for Research Excellence and Technological Enterprise (CREATE), 138602, Singapore

**Keywords:** pupillometry, biofeedback, neurofeedback, arousal, locus coeruleus, noradrenaline, self-regulation, fMRI, oddball

## Abstract

The brain’s arousal state is controlled by several neuromodulatory nuclei known to substantially influence cognition and mental well-being. Here, we investigate whether human participants can gain volitional control of their arousal state using a pupil-based biofeedback approach. Our approach inverts a mechanism suggested by previous literature that links activity of the locus coeruleus (LC), one of the key regulators of central arousal, and pupil dynamics. We show that pupil-based biofeedback enables participants to acquire volitional control of pupil size. Applying pupil self-regulation systematically modulates activity of the LC and other brainstem structures involved in arousal control. Further, it modulates cardiovascular measures such as heart rate, and behavioural and psychophysiological responses during an oddball task. We provide evidence that pupil-based biofeedback makes the brain’s arousal system accessible to volitional control, a finding that has tremendous potential for translation to behavioral and clinical applications across various domains, including stress-related and anxiety disorders.

## Introduction

The brain’s arousal state is controlled by key neuromodulatory nuclei including the noradrenergic (NA) locus coeruleus (LC), dopaminergic substantia nigra/ventral tegmental area (SN/VTA), serotonergic dorsal raphe nucleus (DRN), and the cholinergic nucleus basalis of Meynert (NBM)^1–4^. Previous research indicates that, under constant lighting conditions, pupil size is an indirect ‘indicator’ of the brain’s arousal state. Arousal-related neuromodulatory systems have been directly or indirectly linked to non-luminance related changes in pupil size^5–7^ with the strongest evidence for the LC-NA system^7–11^: Selective chemogenetic or optogenetic activation of the LC causes substantial pupil dilation in mice^11–13^. Further, another animal study suggests the involvement of both cholinergic and noradrenergic systems in pupil dynamics^7^. However, only prolonged pupil dilations during movement were accompanied by sustained cholinergic activity, while moment-to-moment pupil fluctuations during rest closely tracked noradrenergic activity^7^. Furthermore, noradrenergic activity preceded cholinergic activity relative to peak pupil size^7^ and can depolarize cholinergic neurons^1^, suggesting that noradrenergic activity drives cholinergic activation. In humans, functional magnetic resonance imaging (fMRI) demonstrated that LC activity correlates with pupil size, both at rest and during various tasks including the oddball paradigm^6, 8, 9, 14^.

Theories based on intracranial recordings in animals suggest that the LC-NA system modulates functional circuits related to wakefulness, sleep^15–19^ and cognitive processes relevant for task engagement and performance^4, 18, 20–22^. These theories postulate that LC neurons exhibit tonic and phasic discharge patterns, where tonic activity is thought to closely correlate with the brain’s arousal state (i.e., high tonic LC activity is associated with high arousal) and phasic discharge facilitating behavioral responses to task-relevant events^4^. Probing these task-relevant processes with a two-stimulus oddball paradigm revealed that phasic LC responses and task performance depend on the level of tonic activity. For instance, recordings in monkeys showed that elevated tonic LC activity is associated with reduced phasic responses and detection performance of salient oddball stimuli^23–26^.

Regulating arousal is challenging and existing approaches in humans rely mainly on pharmacological agents with side-effects. Here, we investigated an approach utilizing the mechanistic link between the brain’s arousal state and pupil via an innovative pupil-based biofeedback approach (pupil-BF). Only a few previous studies have trained volunteers to self-regulate pupil size with varying degrees of success^27–30^. Volitional pupil size downregulation is especially difficult to acquire^30^. Our main idea is that participants apply different arousing or relaxing mental strategies while receiving online pupil-BF (Fig. 1A). Considering the strong link between pupil diameter and LC-NA activity, we derived several hypotheses from the rodent literature and tested whether pupil self-regulation will affect specific aspects of neural processing^20, 31^ and cardiovascular function^32^ which is influenced by the LC through its projections to autonomic control structures in the brainstem and spinal cord^18^. Specifically, we hypothesized that (i) pupil-BF allows participants to discover suitable mental strategies for volitional up-versus downregulation of pupil size, such self-regulation (ii) is associated with up-versus downregulating activity in brain regions involved in arousal control including the LC and (iii) causes systematic changes in cardiovascular parameters. To further probe the link to the LC-NA system shown to be associated with behavioral and psychophysiological measures of the oddball task, we combined our pupil-BF approach with an auditory version of this paradigm. We hypothesized that (iv) self-regulating pupil size modulates stimulus detection behavior and pupil dilation responses to oddball stimuli.

**Figure 1.**
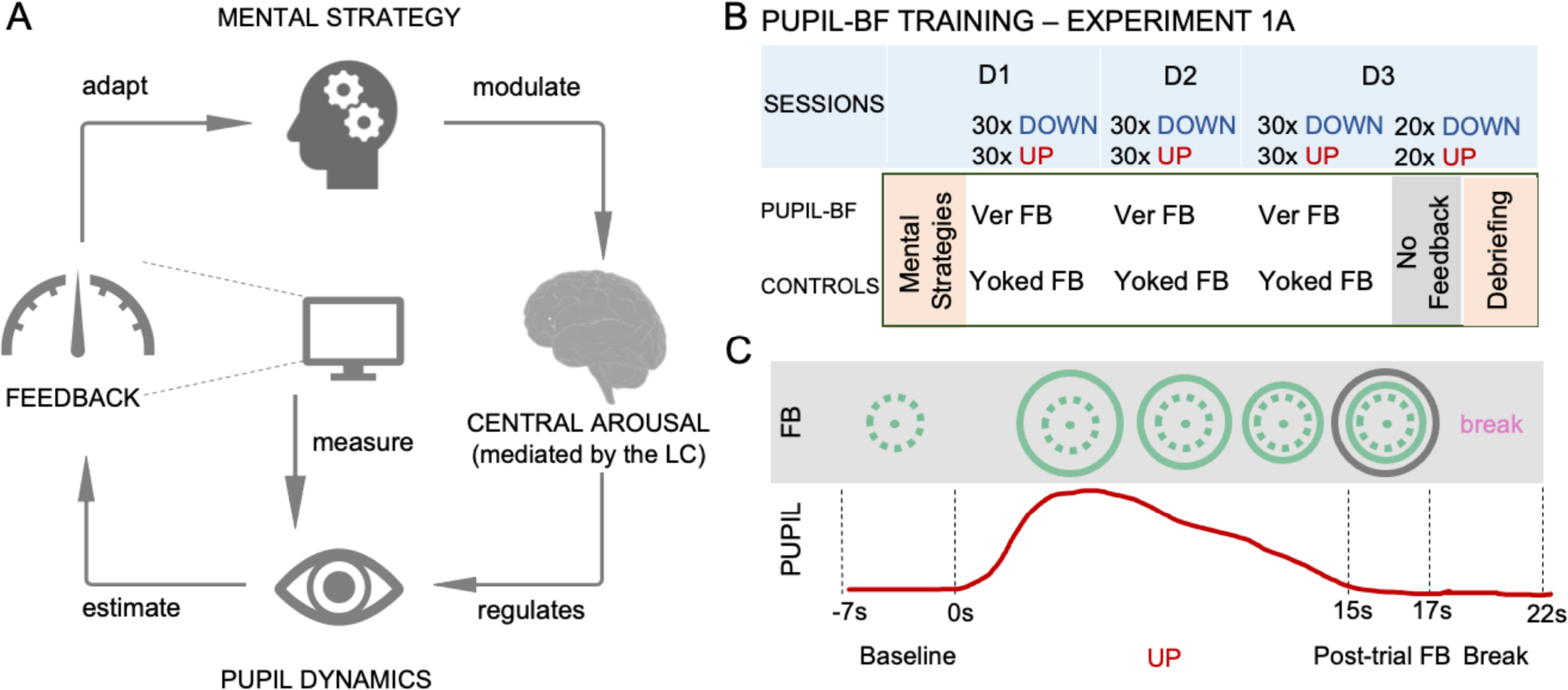
Pupil-based biofeedback (pupil-BF) training. **A.** Participants apply mental strategies which are believed to modulate the brain’s arousal levels mediated by nuclei such as the LC. Pupil size is measured by an eye tracker and fed back to the participant via an isoluminant visual display **B.** In Experiment 1A, healthy volunteers were informed about potential mental strategies of arousal regulation and then participated in 3 days (D1, D2, D3) of upregulation (UP) and downregulation (DOWN) training (30 trials each) while receiving either veridical (ver) pupil feedback (pupil-BF group) or visually matched input/yoked feedback (control groups I and II). At the end of day 3, all participants performed 20 UP and 20 DOWN trials without receiving any feedback and were debriefed on which strategies they have used. **C.** Example trial of the experiment. Each trial consisted of (i) 7s baseline measurements, (ii) 15s modulation phase where the pupil-BF group sees a circle that dynamically changes its diameter as function of pupil size (veridical feedback), (iii) 2s of color-coded post-trial performance feedback (green = average circle size during modulation; black = maximum (UP) or minimum (DOWN) circle size during modulation), and (iv) 5s break. While the upper panel shows an example of what participants would see on their screen, the red line in the lower panel indicates measured pupil size. Note that the control groups I and II see a circle that changes independently of pupil size but resembles that of a participant in the pupil-BF group.

Numerous neuro- or biofeedback studies have shown that providing feedback enables humans to gain remarkable control over specific body functions^33–36^. Acquiring a suitable mental strategy – explicitly or implicitly – is crucial for effective self-regulation and feedback can indeed facilitate this process^33, 36^. Using our pupil-BF training, healthy volunteers learned to volitionally up- and downregulate their pupil size (Experiments 1AB). Crucially, combining pupil self-regulation with fMRI and cardiovascular measurements, we observed systematic modulation of (i) activity in arousal-regulating brainstem centers including the LC and SN/VTA and (interconnected) cortical and subcortical brain regions (Experiment 2), and (ii) heart rate (Experiments 1B,2). Finally, self-regulating pupil size modulated behavioral performance and psychophysiological markers of LC-NA activity during an auditory oddball task (Experiment 3).

## Results

### Experiment 1: Pupil-BF training enables voluntary pupil self-regulation

Participants were randomly assigned to either a pupil-BF (n = 28) or control group (n = 28) and underwent three days of pupil-BF training (Experiment 1A). Prior to training, all participants received instructions on mental strategies derived from previous research^27, 30, 37–39^, such as imagining emotional (i.e., fearful or joyful) situations to upregulate pupil size (UP); or relaxing, safe situations together with focusing on their body and breathing to downregulate pupil size (DOWN; see Supplementary Table (ST)1 for strategies). Each of the three sessions consisted of 30 UP and 30 DOWN trials, with participants self-regulating their pupil for 15s (Fig. 1C). The pupil-BF group received isoluminant visual feedback on pupil size in quasi real-time during regulation and average performance-feedback after regulation (post-trial feedback; Fig. 1C). The control group received feedback from a randomly selected participant of the pupil-BF group (i.e., receiving the same visual input as the pupil-BF group) and participants were instructed to focus on their mental strategies ensuring to control learning effects due to mental rehearsal^40^. Importantly, the control group was not aware of the existence of a pupil-BF group. At the end of day 3, all participants underwent an additional *no feedback-session* (20 UP and 20 DOWN trials) using the same self-regulation strategies as during training, but without receiving feedback.

Descriptively, participants of both groups (n = 27 in each group for final analyses) showed some ability to upregulate pupil size (Fig. 2A), with greater upregulation observed in the pupil-BF group. The ability to downregulate pupil became gradually more successful during training in the pupil-BF group while this ability was generally reduced in the control group (Fig. 2B; Supplementary Figure (SF)1 displays statistical comparisons of up- and downregulation compared to baseline). Importantly, the pupil-BF participants maintained their self-regulation abilities during *no-feedback-trials*, indicating that they acquired a transferable skill that can be applied without constant feedback (Fig. 2AB; grey lines).

**Figure 2.**
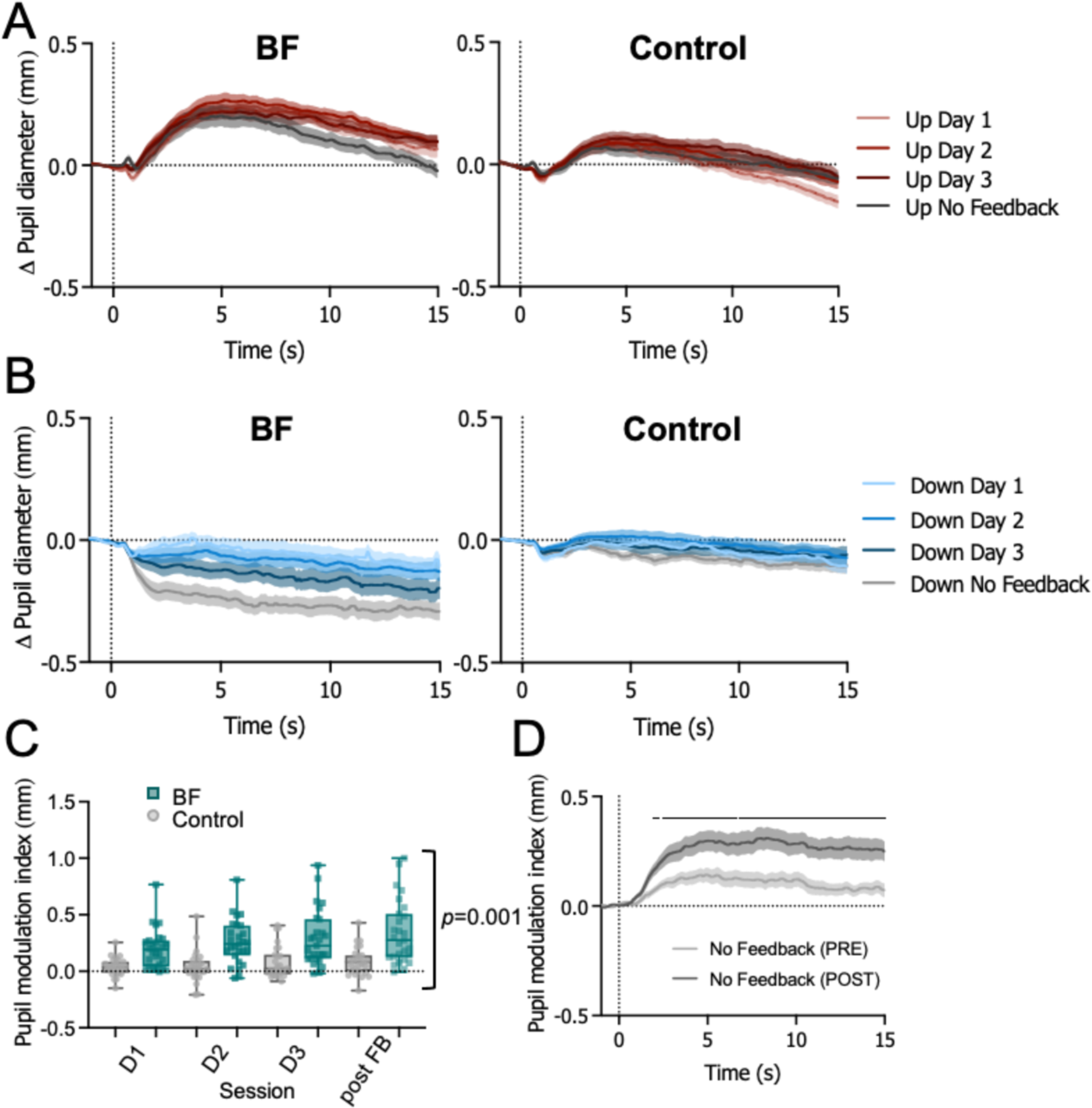
Pupil-BF training results. Average changes in pupil size during 15s upregulation (UP) **(A)** and downregulation (DOWN) **(B)** are shown for the pupil-BF (n = 27) and initial control group (n = 27) for training sessions on Day 1, Day 2, Day 3, and the no feedback post training session of Experiment 1A. **C.** The pupil modulation index reflects the difference between the average pupil size during the two conditions (UP–DOWN) and is shown for each session (Day 1, Day 2, Day 3, no feedback post training session) and group (initial control group versus pupil-BF group of Experiment 1A; dots and squares represent individual participants). Pupil modulation indices were generally higher in the pupil-BF group compared to the initial control group (robust ANOVA, main effect of *group*: *F*(1,21.58) = 21.49; *p* = 0.001,λ_p_^2^ = 0.50; other main effects/interaction *p* ≥ 0.07) **D.** Time series of pupil modulation index measured during the no feedback session before (Pre, light grey) and after pupil-BF training (Post, dark grey) in Experiment 1B (independent cohort, n = 25). Solid black lines at the top indicate clusters of significantly higher modulation indices at Post compared to Pre (SPM1D repeated measures ANOVA; all *p <* 0.05). Shaded areas and error bars indicate SEM. For a replication of results in control group II, see SF 3. For more detailed information on statistical parameters see Supplementary Table (ST) 5.

Pupil self-regulation was quantified calculating a pupil modulation index (UP–DOWN; i.e., the difference between pupil diameter changes in the two conditions) which was significantly larger in the pupil-BF than in the control group (Fig. 2C; *F*(1,21.58) = 21.49; *p* = 0.001,11_p_^2^ = 0.50; 95%-confidence interval (CI)11_p_^2^ [0.17;0.67]). A control analysis confirmed that these group differences were not driven by differences in absolute pupil size at baseline (all *p* ≥ 0.49; SF 2A). These findings indicate that training with veridical BF was superior to mental rehearsal and that the effects were not driven by visual input alone. We tested an additional control group (control II) that received yoked feedback but believed that it was veridical BF (see methods) and thereby controlled for additional motivational and perceived success factors. We obtained similar results as with control group I (SF 3A-C).

Next, we conceptually replicated these results in an independent pupil-BF cohort (Experiment 1B). 26 participants (n = 25 for final analyses) followed a similar 3-day training protocol with a no-feedback-session *before* (Pre) and *after* (Post) training (SF 4AB). Pupil modulation index time series (UP– DOWN) were significantly higher Post as compared to Pre during most of the modulation phase (significant differences: 1.9s – 15s after modulation onset; all *p* < 0.05; Fig. 2D; SF 4E displays condition effects). Interestingly, prior to training (Pre), the modulation index was already significantly higher than 0 (mainly between 1.7s and 10.4s after modulation onset; all *p <* .05; SF 4D), indicating the participants’ ability to voluntarily modulate pupil size only with instructions on mental strategies, an ability that was further improved by pupil-BF training.

In Experiments 1AB, we found that healthy volunteers can learn to self-regulate pupil size. Significant differences in self-regulation capabilities between (i) pupil-BF and control groups and (ii) Pre versus Post performance in a separate pupil-BF cohort highlight the benefits of biofeedback in improving volitional pupil size control.

### Experiment 2: Pupil self-regulation combined with fMRI

Previous studies have repeatedly linked non-luminance-related pupil size changes to activity changes in arousal-regulating centers, including the LC^6, 8–11^. Here, we tested the hypothesis that self-regulating pupil size is associated with activity changes in these regions. 25 trained pupil-BF participants from Experiments 1AB performed pupil up-versus downregulation during two fMRI sessions, one measuring activity across the whole brain, and one specifically measuring brainstem activity (counterbalanced session order). Participants performed 15s of pupil size modulation and received post-trial performance feedback. Pupillometry data confirmed that participants were able to self-regulate pupil size (significant differences between UP and DOWN from 965.72ms (whole-brain, *p* < .001) and 983.72ms (brainstem, *p* < .001) to end of modulation, respectively, Fig. 3A,4A).

**Figure 3.**
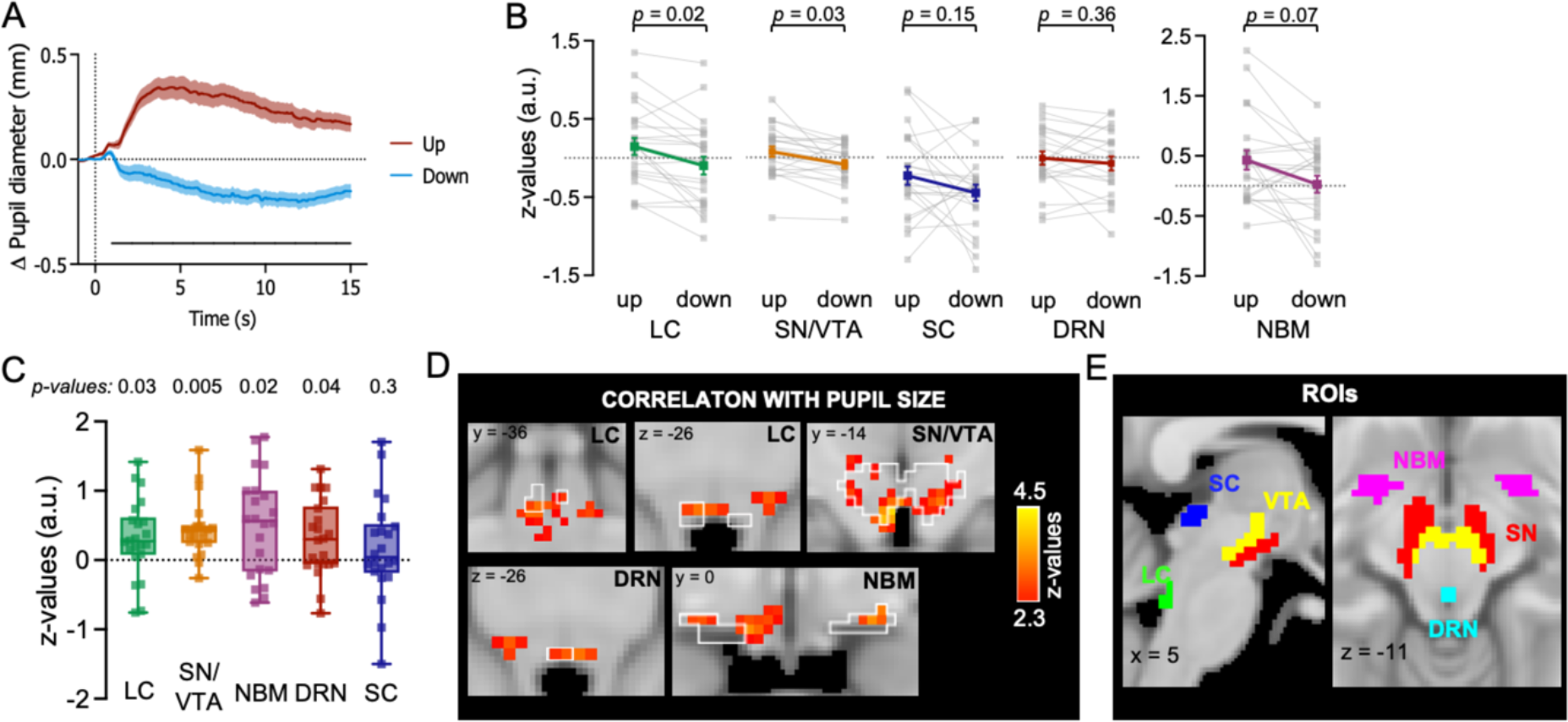
Brainstem fMRI results. **A.** Changes in pupil size averaged across all participants for UP (red) and DOWN (blue) trials showing successful self-regulation of pupil size during brainstem fMRI recording. The solid black line at the bottom indicates a cluster of significantly higher baseline-corrected pupil sizes during UP compared to DOWN (SPM1D paired samples *t*-test; *p <* 0.001; *z*\* = 3.45). **B.** Activity during UP versus DOWN phases of pupil self-regulation in the different regions of interest (ROIs). Statistical comparisons revealed significant effects (UP > DOWN) in the LC and the SN/VTA but not in the SC and DRN. Results for the NBM did not survive correction for multiple comparisons. **C.** Correlation between continuous pupil size changes and BOLD response changes shown as z-values for the different ROIs. Statistical comparison (against 0) revealed significant effects for the LC, SN/VTA, NBM, and DRN. All ROI analyses in **B** and **C** were corrected for multiple comparisons using sequential Bonferroni correction. Squares represent individual data **D. Upper panel:** Correlation analysis revealed that BOLD activity at the LC covaries significantly with continuous changes in pupil diameter (cluster-corrected at z = 2.3; *p* < 0.05). **Lower panel:** Other brainstem areas than the LC exhibited a significant correlation between changes in pupil diameter and BOLD activity (cluster-corrected at z = 2.3; *p* < 0.05). For a complete overview of regions, see ST 2B. The white outlines in **D** indicate different brainstem^43–49^ and basal forebrain regions^98^. **E.** A-priori defined ROIs in the brainstem^43–49^ and basal forebrain^98^ in MNI space. Shaded areas and error bars indicate SEM. Locus coeruleus (LC); substantia nigra (SN) and ventral tegmental area (VTA) combined into one ROI (SN/VTA); dorsal raphe nucleus (DRN); nucleus basalis of Meynert (NBM); and superior colliculus (SC). For more detailed information on statistical parameters, see ST 5.

**Figure 4.**
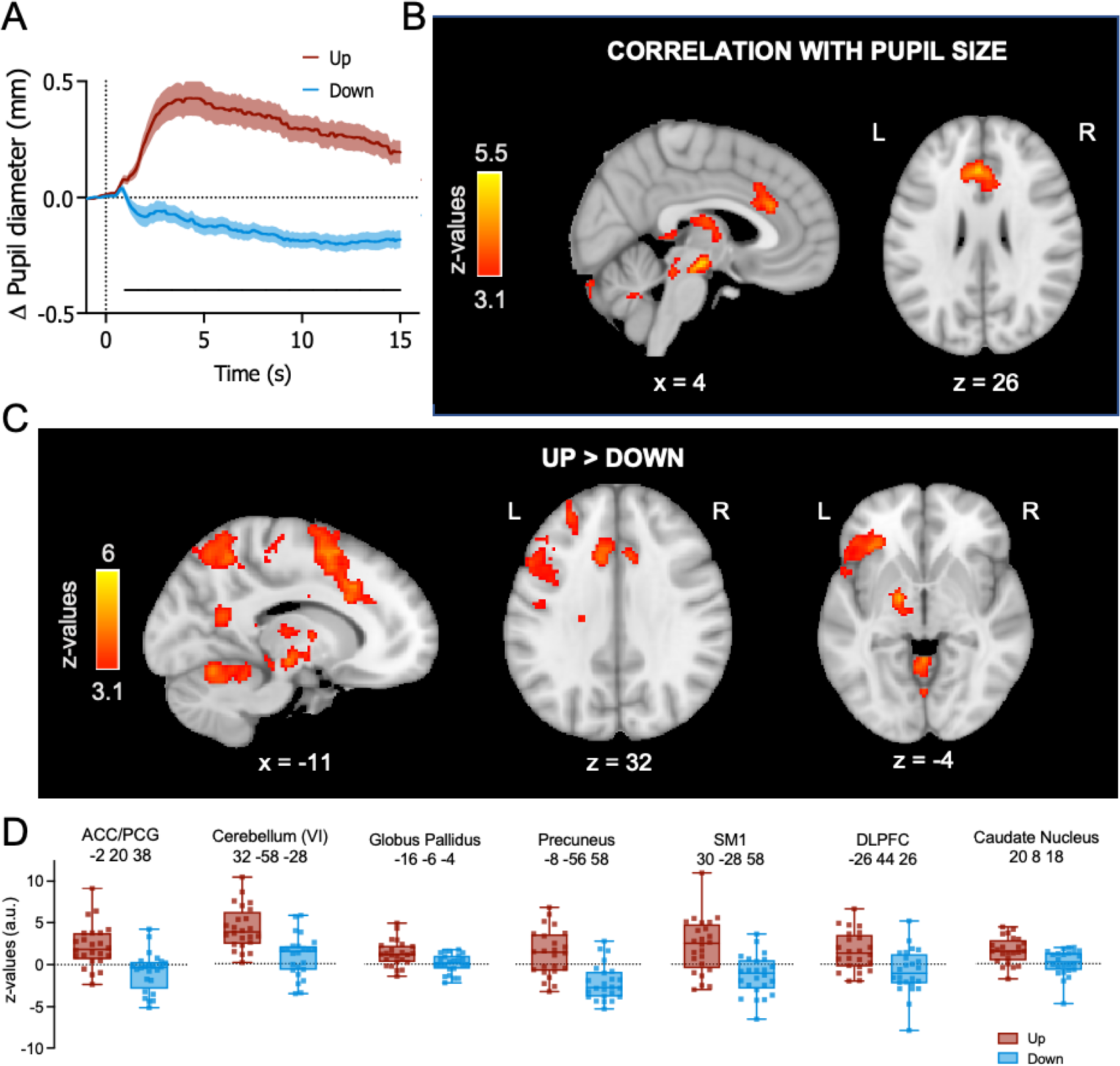
Whole-brain fMRI results. **A.** Changes in pupil size averaged across all participants for UP (red) and DOWN (blue) trials showing successful self-regulation of pupil size during whole-brain fMRI recordings. The solid black line at the bottom indicates a cluster of significantly higher baseline-corrected pupil sizes during UP compared to DOWN (SPM1D paired samples *t*-test; *p <* 0.001; *z*\* = 3.38). **B.** Whole-brain maps showing brain regions where BOLD activity correlates with pupil size changes throughout the fMRI runs **C.** Whole-brain maps depicting brain regions that showed a significant activation during UP as compared to DOWN. All activation maps in **(B)** and **(C)** are thresholded at z > 3.1 and FWE-corrected using a cluster significance level of *p* < 0.05. **D.** Estimated BOLD response represented by z-values for UP vs rest and DOWN vs rest extracted from the peak voxel of each significant cluster shown in C. Squares indicate individual participants. Shaded areas and error bars indicate SEM.

#### Pupil self-regulation is linked to changes in arousal-regulating brainstem regions

Based on previous findings, we predefined the LC, DRN, NBM, and SN/VTA as regions of interest (ROI) that substantially contribute to arousal control^1–4^. Additionally, the LC, NBM and DRN have been linked to pupil size changes^5, 7, 41^ even though it is still debated whether the NBM and DRN modulate pupil size directly or via the LC-NA system. Anatomically, pupil size is controlled by the tone of (i) the dilator pupillae muscle receiving innervation via noradrenergic sympathetic neurons and (ii) the constrictor pupillae muscle receiving innervation via cholinergic parasympathetic neurons^18^. Thus, pupil diameter reflects the relative activity of these opposing outputs of the autonomic nervous system^18, 31^. The LC is assumed to modulate pupil by (i) facilitating sympathetic activity via projections to the intermediolateral (IML) cell column of the spinal cord, and (ii) inhibiting parasympathetic activity via its projections to the Edinger-Westphal nucleus (EWN^18, 31^ reviewed in^41^). Alternatively, a parallel activation of the LC and the sympathetic nervous system through a third player, the rostral ventrolateral medulla, has been discussed in literature^42^. The superior colliculus (SC) is another candidate implicated in non-luminance-related pupil control during the multisensory integration and orienting responses^10, 41^, assumed to influence parasympathetic activity through direct and indirect projections to the EWN via the mesencephalic cuneiform nucleus (MCN), and sympathetic activity through projections to the IML via the MCN^41^. Therefore, we also included the SC as control ROI in our analysis (Fig. 3E).

After pre-processing brainstem data (i.e., removing physiological noise), we extracted mean signal intensities during UP>rest and DOWN>rest from the LC, SN/VTA, DRN, NBM, and SC (n = 22). We observed significantly higher LC activation during pupil upregulation compared to downregulation (*t*(21) *=* 3.40; *p* = 0.015, *d =* 0.73, 95%-CI*d* [0.25; 1.19]). Similar results were found for the SN/VTA (*t*(21) = 2.96, *p* = 0.03, *d =* 0.63, 95%-CI*d* [0.17; 1.08]). Activation differences in all other ROIs did not survive correction for multiple comparisons (NBM: z *=* 2.26, *p* = 0.072, r = 0.48) or did not reach significance (SC and DRN, all *p* ≥ 0.15; Fig. 3B). For additional control analyses extracting mean signal intensities from masks covering (i) the complete brainstem including the midbrain, pons, and medulla oblongata and (ii) the 4^th^ ventricle as well as for all ROI analyses without spatial smoothing of the data, see SF 5A-C and SF 6.

Further correlating pupil size modulations throughout fMRI runs (pupil shifted by 1s) with continuous BOLD timeseries extracted from the pre-defined ROIs revealed a significant association in the LC (*t*(21) *=* 2.64; *p* = 0.03, *d =* 0.56, 95%-CI*d* [0.11; 1.01]), SN/VTA (*t*(21*) =* 5.13; *p* = 0.005, *d =* 1.09, 95%-CI*d* [0.56; 1.62]), NBM (*t*(21) *=* 3.21; *p* = 0.02, *d =* 0.68, 95%-CI*d* [0.21; 1.14]), and DRN (*t*(21) = 2.69; *p* = 0.04, *d =* 0.57, 95%-CI*d* [0.12; 1.02]). SC activity was not significantly related to pupil (0 = 0.32). For additional analyses on unsmoothed data, see SF 6.

Next, complementing our a-priori-defined ROI analysis, we conducted exploratory analyses for UP>DOWN contrasts across the brainstem. However, no region in the brainstem survived cluster-corrections for multiple comparisons (for uncorrected results *p*<0.05, see SF 5D, ST 2A; SF 5E displays DOWN>UP results). Conducting correlations of continuous pupil diameter with continuous BOLD activity changes for every voxel in the brainstem, we found significant correlations in regions covering the SN, VTA, LC, NBM and the DRN (cluster-corrected, Fig. 3D). Additionally, pupil size correlated with activity in other brainstem regions involved in arousal and autonomic regulation ^43–49^(ST 2B).

In summary, our brainstem fMRI analyses revealed that pupil self-regulation is linked to activity changes in brainstem regions involved in arousal and autonomic regulation, including the LC-NA system, with activation during upregulation and deactivation during downregulation. Although the LC is not the only area modulated by pupil self-regulation, our results demonstrate that the reported mechanism of LC activity driving pupil size can be inverted in the context of pupil-BF, making the brain’s arousal system accessible to voluntary control.

#### Pupil self-regulation is linked to changes in cortical and subcortical regions

Investigating the effects of pupil self-regulation on cortical and subcortical structures, we contrasted UP>DOWN phases in our whole-brain fMRI data (n = 24). Up-versus downregulation was associated with significantly higher activation in various brain regions closely connected to the LC, including the dorsal anterior cingulate cortex (dACC)/paracingulate gyrus (PCG), dorsolateral prefrontal cortex (DLPFC), orbitofrontal cortex (OFC), precuneus and thalamus. Additionally, we observed significant activation in primary somatosensory regions (SM1), basal ganglia (globus pallidus, caudate nucleus), and cerebellum (Fig. 4CD; ST 3A).

Examining in which brain areas BOLD activity covaried with pupil size throughout the task, we observed significant effects in the dACC, precuneus, thalamus, globus pallidus, and cerebellum (Fig. 4B; ST 4), partially overlapping with regions activated during pupil size upregulation (UP>DOWN; SF 5FG, ST 3B display DOWN>UP results).

Taken together, pupil self-regulation is linked to an interplay of activation and deactivation in circumscribed brain regions interconnected with the LC, including prefrontal, parietal, thalamic and cerebellar areas.

#### Pupil self-regulation modulates cardiovascular parameters

We tested the influence of self-regulating pupil size on cardiovascular parameters using electrocardiogram (ECG) signals during pupil-BF training (recorded in 15 participants, Experiment 1B) and pulse oximetry data during subsequent fMRI sessions (Experiment 2). We observed higher heart rates during UP than DOWN trials (Fig. 5AC). This difference became more pronounced across pupil-BF training (*condition*\**session* interaction; *F*(2.34, 30.37) = 3.37, *p =* 0.04, 11_p2_ = 0.21, 95%-CI11_p2_ [0.00;0.39]) and remained stable during fMRI (main effect *condition*; *F*(1,22) = 72.25, *p <* 0.001, 11_p2_ = 0.77, 95%-CI11_p2_ [0.54;0.85]; Fig. 5AC). Prior to training, pupil diameter and heart rate changes were largely unrelated (*p* = 0.86, Fig. 5E, left panel). However, during fMRI a higher pupil modulation index (UP–DOWN), thus better pupil self-regulation, was associated with larger differences in heart rate (UP–DOWN; rho = 0.63; *p* = 0.002, 95%-CIrho [0.30;0.82]; sequential Bonferroni-corrected; Fig. 5E, right panel).

**Figure 5.**
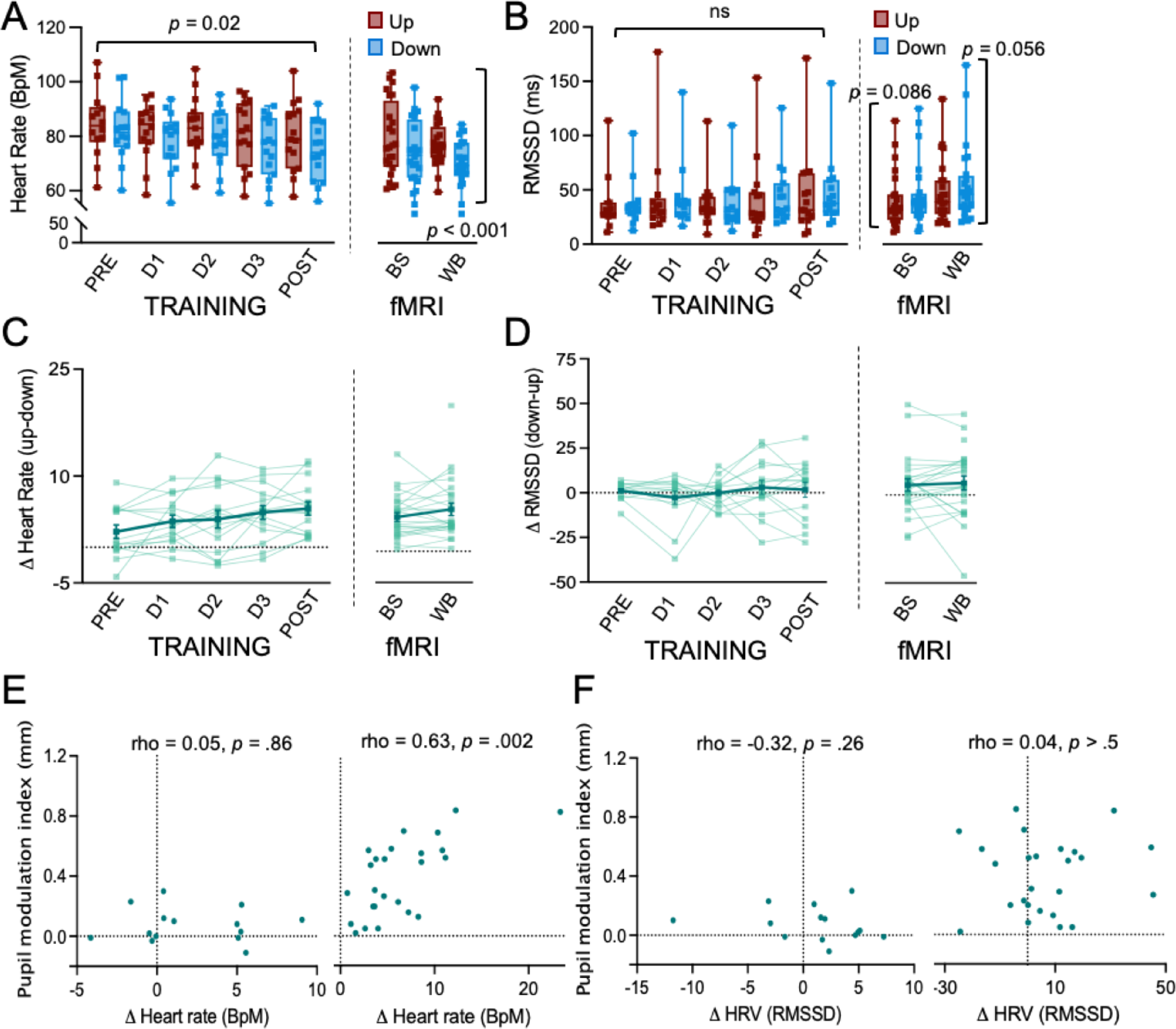
Effects of pupil self-regulation on cardiovascular parameters. Heart rate **(A)** and heart rate variability (HRV) **(B)** averaged for UP and DOWN trials across all participants for pupil-BF training (left panel) and fMRI sessions (right panel). HRV was estimated as the root mean square of successive differences (RMSSD). Self-regulation of pupil size systematically modulated heart rate with an increasingly larger difference between UP and DOWN trials over training sessions (repeated measures ANOVA: *condition*\**session* interaction: *F*(2.35, 32.89) = 3.98, *p =* 0.023, 11p^2^ = 0.22; Greenhouse-Geisser-corrected) which remained stable after training during fMRI (repeated measures ANOVA; main effect *condition; F*(1,22) = 72.25, *p <* 0.001, 11p^2^ = 0.77). Self-regulation did not significantly modulate HRV during training (robust ANOVA: *p >* 0.34). After training during fMRI, HRV was descriptively higher during DOWN than to UP, however, statistical comparisons did not reach significance (Wilcoxon Signed Rank Test; brainstem session: *p* = 0.086; whole-brain session: *p* = 0.056). **C.** and **D.** show individual differences in heart rate (UP-DOWN differences) and RMSSD (DOWN-UP differences) for training (left panels) and fMRI sessions (right panel). The thick solid line represents the group average, thin lines represent individual data. **E.** Spearman rho correlation coefficients between pupil modulation indices (i.e., the difference between pupil diameter changes in the two conditions; UP-DOWN) and differences in heart rate (UP-DOWN) revealing a significant link following (right panel; during fMRI) but not prior to pupil-BF training (left panel) **F.** Non-significant Spearman rho correlation coefficients between pupil modulation indices (UP-DOWN) and RMSSD differences (DOWN-UP) prior to (left panel) and after pupil-BF training during fMRI (right panel). Error bars indicate SEM. BS = brainstem fMRI; WB = whole-brain fMRI. For more detailed information on statistical parameters, see ST 5.

To assess heart rate variability (HRV) measures suggested to represent a marker of parasympathetic activity^50, 51^, we calculated the root mean square of successive differences (RMSSD) and the percentage of successive cardiac interbeat intervals exceeding 35ms (pNN35). There was no clear pupil self-regulation training effect (all *p* ≥ 0.33) on RMSSD (left panel, Fig. 5BD) and pNN35 (SF 7A) and pupil diameter changes were not significantly related to RMSSD (Fig. 5F) and pNN35 (SF 7B) changes prior to and following training during fMRI (all *p* ≥ 0.19). However, pNN35 values were generally higher during down-than upregulation, both during training (z = 2.84; *p* = 0.005; r = 0.76) and subsequent fMRI sessions (whole-brain: z = 3.06; *p* = 0.002; r = 0.62; brainstem: z = 3.371; *p* = 0.002; r = 0.69; sequential Bonferroni corrected, SF 7A). Similarly, during fMRI, RMSSD was descriptively higher during pupil down-than upregulation, however, effects did not reach significance (brainstem: z = 1.71, *p =* 0.086, r = 0.35; whole-brain: z = 1.91; *p =* 0.056, r = 0.39; right panel, Fig. 5BD). We further showed that (learning) effects on heart rate or HRV were weaker or absent when participants trained without BF (control group II; SF 3DE). However, correlations between heart rate and pupil modulation indices revealed similar links as in the pupil-BF group (SF 3F).

Taken together, pupil self-regulation modulated cardiovascular parameters, particularly heart rate, consistent with the model of LC’s role in autonomic function.

### Experiment 3: Combining pupil self-regulation with an auditory oddball task

To determine whether pupil self-regulation modulates behavioral and psychophysiological measures previously linked to LC-NA activity, we combined our pupil-BF approach with an auditory oddball task (Fig. 6A). 22 participants that underwent pupil-BF training (Experiments 1AB) performed the task while (i) upregulating pupil size (ii) downregulating pupil size or (iii) executing a cognitive control task of silently counting backwards in steps of seven. The control task was included to control for cognitive effort effects that may arise from simultaneously executing pupil-self regulation and the oddball task.

**Figure 6.**
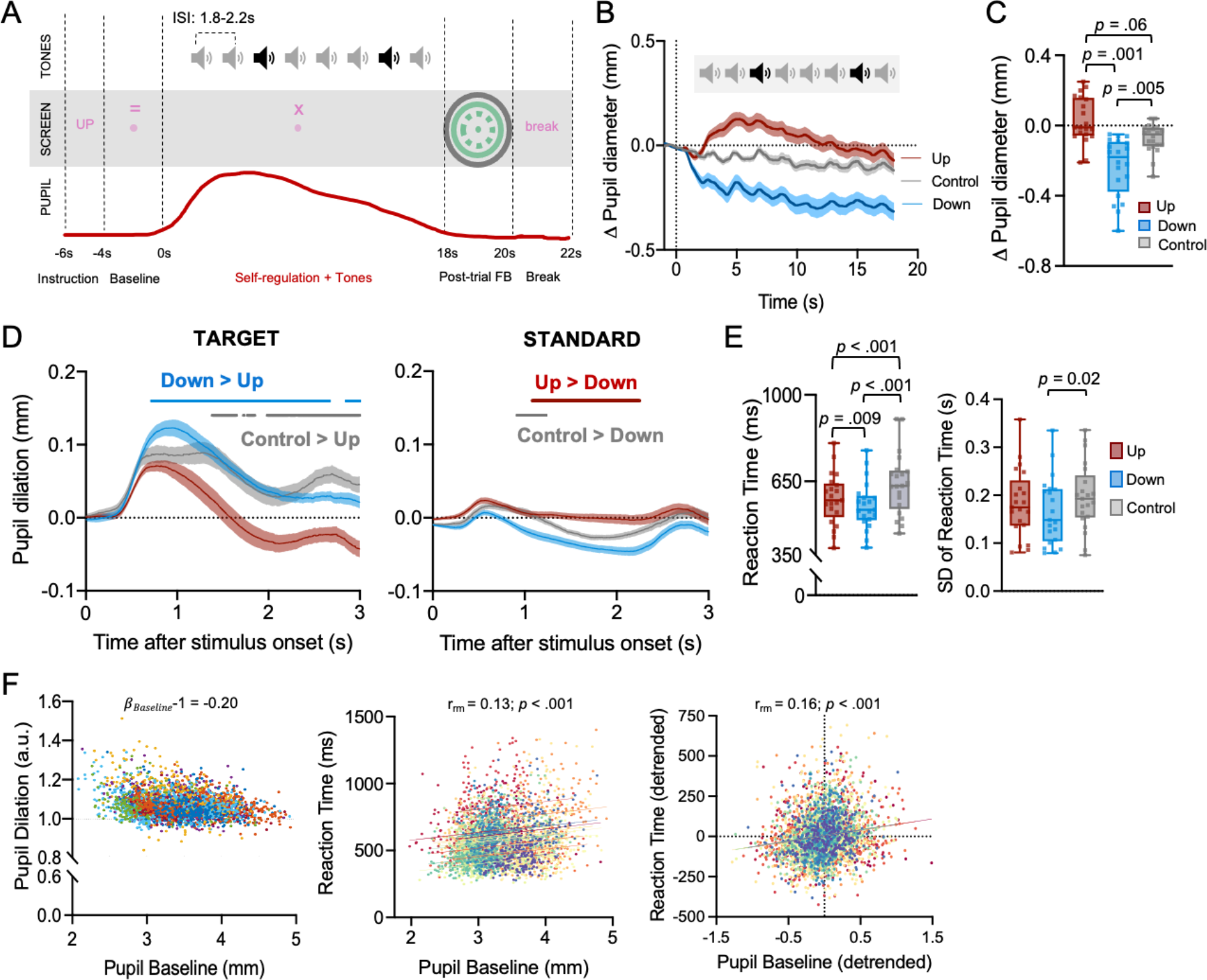
Self-regulation of pupil size combined with an auditory oddball task. **A.** Schematic depiction of an example trial (UP) of Experiment 3 combining pupil-BF and the oddball task in which participants reacted to target tones (black sound icon) by button press and ignored standard tones (grey sound icon) while simultaneously up- or downregulating pupil size or counting backwards (i.e., control). **B.** Pupil size changes averaged across participants for UP (red), DOWN (blue) and control trials (grey, counting backwards in steps of 7) showing the last second of the baseline phase as well as the complete modulation phase of 18s for 20 participants. **C.** Pupil size changes during the modulation phase from baseline (last 1s of baseline phase) averaged across the respective condition showing significantly lower values in DOWN as compared to control and UP trials (robust repeated measures ANOVA with the factor *condition: F*(1.52,16.67) = 9.33, *p* = 0.003, 11p^2^ = 046; DOWN vs. UP: 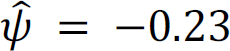; *p* = 0.001; DOWN vs. control: 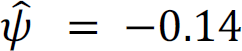; *p* = 0.005; for UP vs. control: 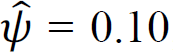; *p =* .06). **D.** Baseline-corrected pupil dilation responses evoked by target (left) and standard tones (right) for UP, DOWN and control trials. Solid lines above the dilation time series indicate time windows of significantly smaller responses to targets in UP as compared to DOWN (blue) and control trials (grey; left panel) and significantly larger responses to standard tones in UP (red) and control (grey) than in DOWN trials (right panel; all tests are Bonferroni-corrected post-hoc tests of SPM1D repeated measures ANOVA with the factor *condition;* all *p <* 0.05) **E.** Behavioral performance of 21 participants during the oddball task depicting faster responses during DOWN as compared to UP (repeated measures ANOVA main effect *condition*: *F*(2,40) = 35.97, *p <* 0.001, 11p^2^ = 0.64, DOWN versus UP: *t*(20) = -2.87, p = 0.009, d = 0.63) and control trials (DOWN versus control: *t(*20) = -7.19, *p* < 0.001, d = 1.57; UP versus control: *t*(20) = -6.04, *p* < 0.001, d = 1.32; left panel; all sequential Bonferroni-corrected). Responses were also less variable (right panel) in DOWN as compared to control trials (*t*(20) = -3.01, *p =* 0.02, d = 0.66; sequential Bonferroni-corrected post-hoc tests of repeated measures ANOVA on reaction time and SD of reaction times with the factor *condition*). **F.** Results of single-trial analyses linking baseline pupil size (500ms prior to target onset) with relative pupil dilation responses (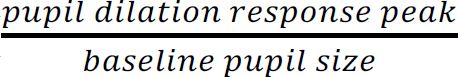; left panel) and reaction times (middle panel: raw values; right panel: detrended values) towards targets. Please note that sound icons in **A.** and **B.** indicate standards (grey) and targets (black) played during pupil self-regulation or control trials. Squares in **C.** and **E.** represent individual data. Error bars and shaded areas indicate SEM. For more detailed information on statistical parameters, see ST 5.

Participants (n = 20 for final analyses) successfully self-regulated pupil size even when performing the oddball task simultaneously (main effect *condition*: *F*(1.52,16.67) = 9.33, *p* = 0.003, 11_p2_ = 0.46, 95%-CI11_p2_ [0.08; 0.65]; Fig. 6BC) as indicated by smaller (baseline-corrected) pupil sizes during DOWN than UP and control trials. Also, absolute pupil size was increased for UP as compared to DOWN trials only during self-regulation but not during the baseline phase (SF 2C,8).

Unexpectedly, absolute pupil size was increased for control trials (SF 2C,8). Since this increase was observed during the baseline *and* self-regulation phase, baseline-corrected pupil sizes were at intermediate levels compared to UP and DOWN conditions (Fig. 6B). This suggests a potentially higher cognitive load during counting than pupil self-regulation throughout all phases of this control condition.

#### Pupil self-regulation modulates psychophysiological markers of LC activity

Previous research in monkeys suggests that elevated tonic LC activity constrains the intensity of phasic LC responses to salient stimuli^24–26^. Here, we tested the prediction that phasic activity varies depending on tonic activity levels. We particularly investigated whether sustained pupil self-regulation is associated with changes in pupil dilation responses to target sounds during the oddball task, which has been linked to phasic LC activity. Consistent with previous research, we observed that pupil self-regulation led to differences in pupil dilation responses to target sounds (*z* =* 7.29; 714 to 3000ms, *p <* 0.001, Fig. 6D, left panel) with significantly larger initial and prolonged elevated pupil dilation during DOWN compared to UP trials (*z\** = 4.07; 718 to 2669ms, *p <* 0.001; 2851 to 3000ms, *p* = 0.005, Bonferroni-corrected).

The cognitive control and UP conditions caused a similar initial pupil dilation but their time courses after peak dilation differed significantly (*z\** = 4.12; between 1381 and 3000ms, all *p* σ; 0.02; Bonferroni-corrected, Fig. 6D, left panel). Differences between control and DOWN trials were not significant. In an exploratory analysis, we examined the single-trial relationship between baseline pupil size and pupil dilation. We found that smaller baseline pupil sizes before target onset were significantly related to larger pupil dilation responses towards target sounds (***β****_Baseline_* - 1 = -0.20, *t*(19) *=* -10.42, Cohen’s *d* = -2.33, 95%-CI*d* [-3.18; -1.47]; Fig. 6F, left panel; SF 9A displays results within conditions).

Standard tones evoked only a minor pupil response (Fig. 6D, right panel). Significant differences in responses between conditions were mainly driven by sustained elevation of pupil size in UP compared to DOWN trials (*z\** = 3.93: 1081 to 2251ms, *p <* 0.001; Bonferroni-corrected) and a faster decrease in pupil size in DOWN compared to control trials (*z\** = 4.23, between 916 and 1233ms, all *p* :: 0.02; Bonferroni-corrected). However, these differences were observed *after* peak dilation and likely reflect the effects of pupil self-regulation.

#### Pupil self-regulation modulates oddball task performance

LC activity has been linked to the detection of task-relevant target stimuli^8, 23, 25, 52^. Here, we examine whether pupil self-regulation, which modulates LC activity, influences oddball task performance. Overall, accuracy was high (Control: 94.3%, UP: 95.8%, DOWN: 97.9%). Analyzing reaction times to target sounds, we found that participants responded slowest during cognitive control and fastest during DOWN trials (Fig. 6E, left panel, main effect *condition*: *F*(2,40) = 35.97, *p <* 0.001, 11_p2_ = 0.64; 95%-CI11_p2_ [0.43; 0.74]). Evaluating the relationship between single-trial pre-target baseline pupil size and reaction times to targets revealed a significant but only weak positive correlation, indicating that smaller pupil size at target onset may be associated with faster responses at single-trial level (r_rm_ = 0.13, *p <* 0.001, 95%-CIr_rm_ [0.11; 0.17]; Figure 6F, middle panel). Considering that both baseline pupil size and reaction times were influenced by a time-on-task effect (linear mixed effects models on reaction times: estimate: 0.00039; t = 7.97; *p* < .001; on pupil size: estimate: -0.00075; t = -6.95; *p* < .001) with *decreases* in pupil size and *increases* in reaction time with time spent on task, we detrended both variables and repeated the analysis revealing a slightly stronger positive correlation (r_rm_ = 0.16, *p <* 0.001, 95%-CIr_rm_ [0.13; 0.20]; Fig. 6F, right panel; SF 9BC display correlations within conditions). Finally, task performance variability measured as the standard deviation (SD) of reaction times, differed significantly between conditions (main effect *condition: F*(2,40) = 4.84, *p =* 0.01, 11_p2_ = 0.19, 95%-CI11_p2_ [0.01; 0.37]); Fig. 6E, right panel) mainly driven by less variable responses in DOWN than control trials.

In summary, self-regulating pupil diameter as a proxy of (LC-mediated) arousal influences behavioral responses as predicted by current theories of noradrenergic function.

## Discussion

In a series of experiments, we showed that participants gain volitional control over their brain’s arousal state via a pupil-BF approach based on the previously suggested mechanistic link between LC-NA activity, arousal and pupil size. We found that healthy adults can learn to self-regulate pupil size during a 3-days training. This ability was significantly reduced when receiving no veridical feedback (Fig. 2, SF 3, 4). Investigating the neural, physiological, and behavioral consequences of up-versus downregulated pupil size revealed three main findings: First, pupil self-regulation significantly modulates activity in arousal-regulating centers in the brainstem, including LC and SN/VTA (Fig. 3). Second, consistent with findings showing that the LC exerts a strong influence on the cardiovascular system^18, 32, 53^, we observed systematic changes in cardiovascular parameters, particularly in heart rate (Fig. 5). Third, pupil-self-regulation significantly influenced task performance and a psychophysiological readout of phasic LC activity during an oddball task (Fig. 6).

Previous research demonstrated the feasibility of achieving volitional control over body and brain functions through bio- or neurofeedback combined with suitable mental strategies in an appropriate task setting^33, 35, 36, 54^. We showed that feedback on pupil diameter significantly improved the ability to volitionally up-versus downregulate pupil size when compared (i) within participants from pre-to post-training and (ii) between veridical pupil-BF and control groups. Upregulation was strong already during the first biofeedback session, while downregulation gradually improved over training. This is in line with previous reports^28, 30^, where upregulating but not downregulating pupil diameter was successful when participants received one biofeedback-training session^30^. Further, once acquired and optimized, pupil self-regulation became a feedforward-controlled skill independent of constant feedback. We tested this ability in a no-feedback-phase immediately after training, however, previous studies showed that self-regulation of central nervous system activity can last beyond the training period indicating that this skill can be retained over time^33^. One concern considering pupil measurements during real-time feedback relates to screen and perceived color luminance. Although we matched perceived color luminance to the grey background of the screen, we cannot rule out inter-individual differences in perceptual luminance. Therefore, the number of colored pixels on the screen was kept constant throughout the feedback phase. Furthermore, our replication experiment without visual interference and only post-trial feedback on the last training day confirmed stable pupil self-regulation even in the absence of online feedback. Another potential concern is the lack of double-blinding in Experiment 1A, since the experimenter knew whether participants belonged to control or pupil-BF groups. However, throughout training, participants were physically isolated from the experimenter in a shielded room. Also, data pre-processing was conducted in an automated manner without knowledge of condition (and group) assignments.

Our hypothesis that volitional pupil size modulation is linked to activity changes in brain nuclei regulating the brain’s arousal level including the LC was confirmed by our ROI analysis on brainstem fMRI data showing that pupil size up-versus downregulation was indeed associated with systematic LC BOLD activity changes. Since neuromodulatory systems don’t act in isolations, self-regulating pupil diameter did not exclusively modulate LC activity but led to analogous activity changes in the dopaminergic SN/VTA and less consistently in the cholinergic NBM. Interestingly, only activation differences in the LC but not in the SN/VTA or NBM significantly exceeded general activation changes across the whole brainstem (SF 5C). Our results, especially regarding LC, were confirmed in additional ROI control analyses without the application of spatial smoothing which led to comparable result patterns. These findings are generally in line with a recent study in humans that linked noradrenergic, cholinergic, and dopaminergic activity to pupil responses during a cognitive task^6^. The co-activation of the adrenergic and dopaminergic system is not surprising since previous studies in non-human primates and rodents have identified noradrenergic LC projections to the VTA^55, 56^ and SN^55^. Based on our methodology, however, we cannot differentiate whether SN/VTA modulation during pupil self-regulation occurs directly through top-down control or indirectly via the LC.

In our study, links between up-versus downregulating pupil size and the NBM did not survive multiple comparison correction and were less consistent than for dopaminergic and noradrenergic regions. Previous comparisons of cholinergic and noradrenergic systems and their role in pupil dynamics in mice provided correlative evidence for the involvement of both systems. However, sustained cholinergic activity was observed mainly during longer-lasting pupil dilations, such as during locomotion, while moment-to-moment pupil fluctuations during rest closely tracked noradrenergic activity^7^. Furthermore, noradrenergic activity preceded cholinergic activity relative to peak pupil dilation^7^, suggesting together with the finding that noradrenergic neurons can depolarize cholinergic neurons^1^, that noradrenergic activity may drive cholinergic activation. However, temporal resolution of fMRI measures is insufficient to reliably test this proposal in our human dataset.

ROI analyses revealed that up-versus downregulating pupil diameter did not systematically modulate BOLD responses in the DRN and SC, brainstem regions implicated in non-luminance-dependent pupil size changes^5, 41^. The absence of an effect in the SC aligns with the theory that it modulates pupil mainly in the context of specific orienting responses towards salient events while LC modulates pupil in the context of arousal^57, 58^.

Explorative general linear model analyses (GLM) on brainstem fMRI and its covariation with pupil size throughout the experiment revealed activation in the LC, SN/VTA, NBM, and DRN but also in other critical nodes for producing a waking state and regulating autonomic activity, including the pedunculopontine nucleus and periaqueductal gray (ST 2B). Even though GLM analyses contrasting up-versus downregulation trials did not survive cluster-corrections, they revealed qualitatively similar activation patterns (including the pre-defined ROIs; SF 4D, ST 2A). This emphasizes that pupil self-regulation may modulate a distributed brainstem network associated with arousal regulation. However, it remains unclear whether these effects are directly driven by cortical and/or subcortical top-down control mechanisms or mediated through the LC as a major relay station. Future research could combine pupil-BF with pharmacological agents targeting different systems to unravel whether the LC is orchestrating the seemingly synchronized activity changes of different neuromodulatory systems and brainstem nuclei during pupil self-regulation.

Importantly, imaging the human brainstem, particularly the LC, is challenging due to the small-sized nuclei, high susceptibility to physiological noise, and lower signal-to-noise ratio than cortical signals. We obtained consistent results across different smoothing levels and additionally applied stringent noise control through independent component analysis (ICA) and physiological noise modeling (PNM) which considers heart rate as a nuisance regressor. Given that up-versus down-regulation was associated with significant differences in heart rate, the fMRI analyses revealed BOLD changes over and above this heart rate effect, indicating that our analysis revealed a conservative estimate of how pupil self-regulation modulates the activity of arousal-related brainstem nuclei.

Our study identified the ACC, OFC, DLPFC, precuneus, thalamus, globus pallidus, and cerebellum as candidate areas that might exert top-down control of the arousal system in the brainstem. All these brain regions are heavily interconnected with the brainstem and particularly the LC. It is tempting to speculate that frontal areas like the ACC and OFC that have dense projections to the LC in non-human primates^4, 59^ form the brain’s intrinsic control system of arousal and exert top-down control of LC activity. However, the nature of this activity, whether it is causal or consequential to arousal modulation, cannot be determined from our data.

Together, our fMRI results demonstrate that pupil size may provide an active information channel for self-regulating activity in areas involved in arousal regulation. Our results implicate the LC as one of the brainstem areas that are significantly modulated by pupil self-regulation, potentially influencing downstream areas involved in arousal control.

The oddball task has been closely linked to LC-NA activity in animal models and human research and pupil dilation responses evoked by oddball stimuli have been considered a psychophysiological marker of phasic LC activity^4, 60, 61^. Additionally, well-known theories derived from work in animal models postulate that phasic LC activity in response to task-relevant events depends on tonic LC activity: when tonic activity is upregulated, phasic responses are weak; whereas when tonic activity is relatively lower at an intermediate level, phasic responses are strong^4, 26^. A similar relationship has been observed in human pupil measurements reporting an inverse relationship between naturally fluctuating baseline pupil size and pupil dilation responses^60–62^. Consistently, we found that downregulating pupil diameter in a sustained way led to larger pupil dilation responses to target sounds. By contrast, upregulating pupil diameter led to smaller pupil dilation responses. These results are consistent with previous work, suggesting that self-regulating pupil diameter modulates tonic LC activity. One concern is that mechanisms specific to the structure of the eye or pupil’s musculature might have limited pupil responses when baseline pupil diameter was already high. However, this is unlikely as previous research has shown that varying pupil diameter through different luminance conditions did not affect task-evoked pupil dilation responses^60, 63^. Accordingly, the more likely conclusion is that pupil dilation responses depend on the brain’s arousal state as reflected in baseline pupil size.

Our behavioral findings that task performance was better when baseline pupil diameter was low during downregulation than when it was high during upregulation or counting require careful interpretation. First, our design does not allow to determine whether pupil downregulation enhances behavioral performance compared to no dual task. This should be addressed in future studies implementing a resting control condition. Second, although our findings align with some results of previous studies analyzing spontaneous^60^ or experimentally induced pupil size fluctuations^64^, there are studies reporting opposite effects in sustained attention tasks^65, 66^. This inconsistency may be attributed to a suggested inverted-U relationship between arousal levels and task performance^4^: tasks that are naturally “non-arousing”, e.g. due to few external stimuli or low cognitive demands might benefit from upregulation, while tasks with more arousing properties, e.g. processing frequent external stimuli or dual-task-conditions, might benefit from downregulation^67^. Overall, our data supports the idea that attentional performance is influenced by baseline pupil size prior to target onset, reflecting arousal levels and potentially tonic LC activity. This influence may be achieved through modulation of cortical processes through arousal-regulating brainstem nuclei including the LC that facilitate adequate behavioral responses.

In our oddball task, we included a control condition where participants count backwards in steps of seven while responding to target tones. Surprisingly, absolute pupil size was substantially higher in this condition already at baseline suggesting increased cognitive effort throughout this task. Despite the differences in absolute pupil size, there was no significant difference in pupil dilation responses to target sounds between the control and downregulation conditions. Furthermore, participants exhibited the slowest and most variable responses to task-relevant sounds in control trials, possibly due to dual-task costs of counting and responding to target sounds. These costs may have been reduced in self-regulation trials as participants had practiced this skill over several days, resulting in more automated processes. This interpretation is consistent with previous EEG-neurofeedback findings comparing task performance during veridical neurofeedback versus a control condition^67^.

Since the LC plays a role in controlling autonomic activity through projections to cardiovascular regulatory structures^18, 32, 53^, we investigated whether pupil self-regulation affects cardiovascular parameters. Consistent with our hypothesis, heart rate was generally higher during pupil up-than downregulation, an effect that increased across training sessions and correlated with the ability to self-regulate pupil size after pupil-BF training. However, in our second control group in which we recorded ECG data throughout training, too, we saw a similarly strong link between pupil self-regulation and differences in heart rate at the end of training. Thus, whether this established link may be due to feedback training or rather linked to the repeated exposure to explicit mental strategies needs to be clarified in future studies. Further, the effects on HRV were less clear. Whereas RMSSD did not significantly differ between pupil up- and downregulation we observed significant effects for the pNN35 (SF 7A). However, it’s worth noting that our pupil self-regulation period was only 15s which is rather short for determining HRV changes. Extending pupil self-regulation duration will enable us to investigate whether this intervention can modulate HRV over longer time scales.

In summary, our study demonstrates that our pupil-BF approach enables healthy volunteers to volitionally control their pupil size. Self-regulation of pupil size is associated with systematic activity changes in brainstem nuclei that control the brain’s arousal state, including the LC and SN/VTA. Moreover, we observed that self-regulation of pupil size modulates (i) cardiovascular parameters and (ii) psychophysiological and behavioral outcomes of an oddball task previously linked to LC activity. Our pupil-BF approach may constitute an innovative tool to experimentally modulate arousal-regulating centers in the brainstem including the LC. Considering the strong modulatory effects of such centers on cognitive function and various behaviors including stress-related responses, pupil-BF has enormous potential to be translated to behavioral and clinical applications across various domains.

## Methods

### General information

All participants included in the study were healthy adults, free of medication acting on the central nervous system, with no neurological and psychiatric disorders and with normal or corrected-to-normal vision by contact lenses. All participants were asked to abstain from caffeine intake on the day of testing. All experimental protocols were approved by the research ethics committee of the canton of Zurich (KEK-ZH 2018-01078) and conducted in accordance with the declaration of Helsinki. Except for fMRI measurements, all studies were conducted in a noise-shielded room (Faraday cage) to allow the participants to focus on their task and to keep lighting settings constant at dim light. All participants provided written informed consent prior to study participation and received monetary compensation. None of the experiments were preregistered.

### Experiment 1A: Pupil-BF training to learn to volitionally regulate pupil size

#### Participants

A priori power analyses based on own pilot data of a single training session with 30 trials (unpublished data) aiming for a power level of 80% resulted in a necessary sample size of 31 participants. As we expected training effects of our multi-session approaches, we recruited 28 participants for the BF (24 ± 5 years, 16 females) and 28 participants for our initial control group (24 ± 5 years; 13 females) for Experiment 1A, exceeding the sample size of other neurofeedback studies^33, 68, 69^. Participants were randomly assigned to the pupil-BF or initial control group. The experimenter was aware of an individual’s group assignment, but all participants received identical, standardized instructions and performed the measurements while sitting alone in a shielded room without any interactions with the experimenter. The processing of the data was done in an automatized and blinded way for all participants together, i.e. without knowing to which group or condition the data belonged. For control group II (see below), we recruited additional 16 participants (25.20 ± 6.67 years, 13 females). The data from control group II (which was collected at a later stage) was analysed separately using the same automatized algorithm which was again blind to the experimental condition. One participant of the pupil-BF group needed to be excluded from final data analyses due to the development of an eye blink-related strategy instead of a mental strategy. One participant of the initial control group dropped out after the first session due to personal reasons. This led to a final sample size of n = 27 for the pupil-BF group and n = 27 participants for control group I, respectively.

#### Pupil-based biofeedback

Participants sat alone in a shielded room in a comfortable chair with their chin placed in a chinrest to ensure a stable head position without putting too much strain on the neck of the participants. We kept the height of the chin rest constant across participants and adjusted the height of the chair to accommodate participants^70^. Their eyes were ∼65cm away from the eye tracker (Tobii TX300, Tobii Technology AG, Stockholm, Sweden) that was positioned below the screen (240B7QPJ, resolution: 1680×1050; Philips, Amsterdam, Netherlands) to allow for optimal eye tracking and measurement of pupil size. They were instructed to look at the fixation dot displayed in the center of the screen. Pupil diameter and eye gaze data of both eyes were sampled at 60 Hz. To ensure that participants did not use eye movement-related strategies (e.g., vergence movements, squinting), we additionally videorecorded the right eye of the participants and visually inspected these videos. At the start of each session, the eye tracker was calibrated using a 5-point calibration.

In all three training sessions, participants received online (we refer to it as “quasi real-time” due to slight delays in feedback related to processing and averaging costs) and post-trial feedback on their pupil modulation performance that was based on estimating pupil size of the dominant eye. We accounted for artifacts caused by eye blinks, physiological and measurement-based noise with the following preprocessing steps: (i) rejection of data samples containing physiologically implausible pupil diameter values ranging outside of a pupil size of 1.5 and 9mm – this step also ensured that blinks, recorded with a value of -1, would not be included in feedback shown to participants; (ii) rejection of physiologically implausible pupil size changes larger than 0.0027mm/s. This previously implemented approach^30, 71^ is based on specifications of a study reporting peak velocity of the pupillary light reflex^72^. Finally, to ensure a smooth feedback display, the last two collected and processed pupil size samples were averaged and displayed^30^. During the pupil self-regulation phase, pupil size was displayed on the screen by means of a moving circle (Fig. 1C) centered around the fixation dot. This moving circle showed pupil size relative to a dashed circle representing the mean pupil size of a 5s baseline phase (i.e., when participants did not self-regulate pupil size). Importantly, the moving circle only indicated pupil size changes in the required direction, e.g., getting larger for upregulation and smaller for downregulation trials. If pupil size didn’t change in the required direction, the circle stayed at the size of the baseline circle. To ensure constant screen luminance levels, the thickness of the circle was adjusted relative to its size so that the numbers of pixels shown on the screen were kept constant throughout the modulation phase. After completion of the modulation phase, post-trial performance feedback was displayed. Here, valid pupil diameter samples were averaged across the modulation phase, the maximum change was extracted and displayed on the screen with average feedback being color-coded: If participants successfully modulated pupil size into the required direction (i.e., pupil size during modulation was larger than baseline pupil size in upregulation trials or smaller than baseline pupil size in downregulation trials), the circle indicating the average change was shown in green (Fig. 1C). If pupil size modulation was not successful (i.e., pupil size during modulation was similar to or smaller than the baseline pupil size in the upregulation trials or bigger than baseline pupil size in downregulation trials), the circle was depicted in red. The maximum change was always indicated in black. This intermittent feedback was displayed for 2s.

Throughout the experiment, we ensured that all used colors were isoluminant to the grey background (RGB [150 150 150]) by calculating relative luminance as a linear combination of red, green, and blue components based on the formula: Y = 0.2126 R + 0.7152 G + 0.0722 B. It follows the idea that blue light contributes the least to perceived luminance while green light contributes the most (https://www.w3.org/Graphics/Color/sRGB). Stimulus presentation throughout the experiment was controlled using the MATLAB-based presentation software Psychtoolbox 3.0.17.

All participants underwent three sessions of pupil-BF on separate days within a period of seven days. The pupil-BF training session took place roughly at the same time of the day to keep circadian influences constant. Prior to pupil-BF training on day 1, participants read an instruction sheet explaining the procedures and providing recommended mental strategies derived from previous publications of dilated or constricted pupil size during different mental states and cognitive or emotional tasks^27, 30, 37–39^. Participants were instructed to rely on these (or similar) mental strategies. Furthermore, we determined the dominant eye of each participant (right eye dominance: n = 22 pupil-BF group; n = 20 control group I; n = 13 control group II) using the Miles test^73^ since the displayed feedback during training was determined by the data recorded of the dominant eye.

In each of the training sessions, three up- and three downregulation blocks were performed, each consisting of ten trials (30 UP/30 DOWN trials per day; Fig. 1B). Each trial (Fig. 1C) started with the display of the direction of modulation (either up- or downregulation) in green for 2s on a grey background followed by a baseline phase of 7s. Participants saw a green fixation dot in the center of a screen surrounded by a dashed green circle during baseline. During this baseline phase, participants were instructed to silently count backwards in their heads in steps of four to bring them into a controlled mental state. Then, a 15s modulation phase indicated by the display of an additional solid green circle started in which participants were asked to use mental strategies to up- or downregulate their own pupil size while receiving online pupil-BF. This modulation phase was followed by post-trial performance feedback for 2s. After a break of 5s indicated by the words “short break” in green, a new trial started. After each block, participants could take a short self-determined break before they continued with the next block.

On training day 3, participants performed additional no-feedback-trials following the same overall structure. However, the start of the modulation phase was indicated by a change from an “=” to an “x” in the same color (green) and same position above the fixation dot. No baseline circle was shown and no feedback, neither online nor post-trial performance feedback was provided.

After the pupil-BF sessions, we conducted a debriefing in which participants reported in their own words which mental strategies they used for up- and downregulation, respectively.

#### Training sessions of control groups

Participants of the initial control group received the same instructions on mental strategies for up- and downregulations and the same amount of training as participants of the pupil-BF group. However, veridical feedback was not provided. To control for visual input on the screen, each participant of the control group was randomly matched to one participant of the pupil-BF group and saw the exact same feedback and received the same visual input as the veridical BF participant by providing a “replay” of the feedback screen of the pupil-BF group. This approach assured not only the same visual input during the feedback phase and the same proportion of positive/negative feedback in all groups. Importantly, participants knew that the visual input was unrelated to their own performance (without the knowledge of a “true feedback group”) to prevent the development of illusory correlations and to exclude learning effects related to mental rehearsal^40^. As in the pupil BF group, control participants were instructed to look at the fixation dot in the center of the screen and to apply the mental strategies introduced to them (or similar).

This control group accounts for learning effects related to repeatedly using mental strategies as well as for visual input, however, it is possible that control participants may lack motivation and perception of success^40^. Therefore, we recruited additional 16 participants for a second control group (control group II), which received the same amount of training and instructions on mental strategies as the two other groups. Again, participants received “yoked feedback”, i.e. they were randomly matched to one participant of the pupil-BF group and saw the exact same feedback and received the same visual input as the veridical BF participant by providing a “replay” of the feedback screen. However, they were told to apply their mental strategies *and* use the biofeedback for optimizing performance.

#### Cardiovascular measurements

For control group II, we additionally recorded cardiac data with a Biopac MP 160 system and the accompanying AcqKnowledge software (Biopac Systems Inc., USA). ECG was recorded continuously and sampled at 1000Hz from two electrodes, one attached to the left lower rib and one under the right collarbone. An electrode attached to the left collarbone was used as a reference. All physiological data was recorded and stored on a PC for off-line analysis. We also recorded continuous respiration data by means of a breathing belt (Biopac Systems Inc., USA) that was affixed to the participant’s chest, however, respiratory data is not reported here.

#### Offline data processing and analysis of pupil data

(Pre-)processing of the pupil data was conducted using Matlab R2018a (MathWorks, Inc., Natick, MA, USA). Recorded pupil size (in mm), gaze data, and video recordings were visually inspected to ensure that participants followed the instructions to look at the fixation dot in the center of the screen during the baseline and modulation phase of the experiment and did not use eye movement-related strategies (e.g., vergence movements or squinting). In case of violations during these phases (i.e., squinting/opening eyes systematically, large eye movements/saccades, i.e., deviations > ∼16° and ∼10° of visual angle on the x- and y-axes, respectively) that can potentially bias the validity of measured pupil size^70^, trials were excluded from further analysis (pupil-BF group = 8.50%; control group I = 8.13%; *Mann-Whitney U =* 341.50, *z =* -0.40; *p* = 0.69, control group II = 9.52%; *Mann-Whitney U* to compare with pupil-BF group: *U =* 234.00, *z =* 0.45; *p* = 0.65). Then, pupil data of both eyes were systematically preprocessed using the guidelines and standardized open source pipeline published by Kret and Sjak-Shie^74^. Invalid pupil diameter samples representing dilation speed outliers and large deviation from trend line pupil size (repeated four times in a multipass approach) were removed using a median absolute deviation (MAD; multiplier in preprocessing set to 12^74^) – an outlier resilient data dispersion metric. Further, temporally isolated samples with a maximum width of 50ms that border a gap larger than 40ms were removed. Then, we generated mean pupil size time series from the two eyes which were used for all further analyses reported (see^74^ on how mean times series can be calculated even with sparsely missing samples of one pupil). The data were resampled with interpolation to 1000Hz and smoothed using a zero-phase low-pass filter with a recommended cutoff frequency of four^74^. In a next step, the resulting data were inspected and trials with more than 30% of missing data points across baseline and modulation phases were excluded. Sessions with more than 50% of missing trials of a participant were excluded from further analysis (n = 1 in the pupil-BF group: participant who developed the eye blink-related strategy). Finally, preprocessed pupil diameter was corrected relative to baseline, by computing mean pupil size of the last 1000ms before the start of the modulation phase of each trial and subtracting this value from each sample of the respective modulation phase, as previously recommended^75^.

In a next step, we averaged the baseline-corrected pupil diameter time series during baseline and modulation phases across the UP and DOWN condition of each session (day 1, day 2, day 3, no feedback post) and participant. Furthermore, for each participant and each session, absolute baseline diameter used for trial-based baseline correction (i.e., averaged across 1000ms before modulation onset) was calculated for each condition (i.e., UP and DOWN). To determine whether it is possible to learn to volitionally modulate pupil size, and to determine whether veridical pupil-BF is essential for successful self-regulation, we calculated a pupil modulation index for all participants of both groups for each session (day 1, day 2, day 3, no feedback post) as the average difference between baseline-corrected pupil values during up-versus downregulation across all n=15000 data points in the 15s interval (i.e. upsampled to 1000Hz)

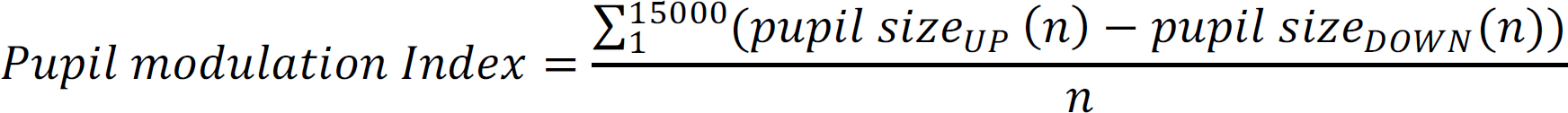

Since successful upregulation is reflected in positive baseline-corrected pupil size values and successful downregulation in negative baseline-corrected pupil size values, larger pupil modulation indices indicate better condition-specific modulation. We further averaged pupil modulation index time series across the complete modulation window to receive one modulation index for each session of each participant for statistical analyses.

Statistical analyses were performed using IBM SPSS 28 (IBM Corporation, Armonk, NY, USA), R version 4.1.2/4.2.2 (R Core Team), and JASP (version 0.16.2; https://jasp-stats.org/). Since Shapiro-Wilk tests revealed significant deviations from Gaussian distribution of some of the residuals of the pupil modulation index (UP – DOWN) for both groups, we computed robust mixed design ANOVAs based on 20% trimmed means using the R package WRS2 (version 1.1-3; https://cran.r-project.org/web/packages/WRS2/index.html) to compare the pupil modulation indices with the within-subjects factor *session* (day 1 versus day 2 versus day 3 versus no feedback post) and the between-subjects factor *group* (pupil-BF versus control I). In case of significant effects, we derived post-hoc *p-* values using bootstrap-based functions implemented in the WRS2 package. To test whether motivational factors or perceived success played an additional role, we repeated the same analysis with control group II (i.e., factor session: day 1 versus day 2 versus day 3 versus no feedback post; factor group: pupil-BF versus control II). Finally, we performed statistical analyses to ensure stable pupil size during baseline phases. To this end, we subjected absolute baseline pupil size averaged for each session and condition across participants of each group to a mixed design ANOVA with the within-subjects factors *session* (day 1 versus day 2 versus day 3 versus no feedback post) and *condition* (UP versus DOWN) and the between-subjects factor *group* (pupil-BF versus control I). Sphericity was assessed using Mauchly’s sphericity test and violations were accounted for with the Greenhouse-Geisser correction. In case of significant effects, we derived post-hoc *p-*values corrected for multiple comparisons using sequential Bonferroni correction^76^. Due to violations from normal distribution of baseline pupil size in control group II, we conducted Wilcoxon-signed rank tests to compare baseline pupil size of up- and downregulation trials of each day. Additionally, we ran a Bayesian ANOVA and Bayesian paired-samples t-tests using the same factors to be able to evaluate whether there is evidence for the null hypothesis (i.e., no difference in baseline pupil size between conditions in neither session nor group) and default priors.

#### Cardiac data (pre-)processing and analyses

ECG R peaks in the second control group were detected automatically and, if necessary, manually corrected using the MATLAB-based toolbox PhysioZoo^77^. Data segments consisting of non-detectable peaks or poor quality were excluded from further analyses. Resulting R-R intervals for which both R peaks were occurring in the modulation phase of the pupil-BF training were extracted and further processed in MATLAB (R2021a). Unfortunately, some of the data was corrupted after data saving (n = 1 for all days, n = 1 for training day 2) or had incomplete trigger information due to technical issues with the trigger box (n = 3 for no feedback trials at day 3) leading to n = 15 data sets on day 1 and day 3, n = 14 data sets on day 2, and n = 12 datasets for no feedback trials on day 2. Heart rate, reflecting cardiovascular dynamics controlled by an interplay between the sympathetic and parasympathetic nervous system, was calculated by dividing 60 through the respective R-R intervals of the modulation phase. Furthermore, we computed the root mean square of successive differences (RMSSD) based on R-R-intervals. We chose RMSSD because it is relatively free from breathing influences^78^ and can be computed for intervals as short as 10s^79, 80^. It represents a beat-to-beat measure of HRV in the time domain that is assumed to give meaningful insights about parasympathetic activity of the autonomic nervous system^50, 51^. Heart rate and HRV (RMSSD) calculated for each modulation phase was averaged across the respective up- and downregulation condition of each training session.

To investigate whether pupil self-regulation without veridical biofeedback systematically modulates cardiovascular parameters, we subjected heart rate averages for each condition (UP versus DOWN) and training session (day 1, day 2, day 3, no feedback post) to a two-way repeated measures ANOVA. Sphericity was assessed using Mauchly’s sphericity test and violations were accounted for with the Greenhouse-Geisser correction. Since our HRV measure, RMSSD, significantly deviated from normal distribution (Shapiro-Wilk tests, all *p <* 0.01), we calculated differences scores in RMSSD between up- and downregulation trials (DOWN-UP) for each session (day 1, day 2, day 3, no feedback post). Positive values indicated larger HRV in DOWN than UP trials and negative values indicated larger HRV in UP than DOWN trials. These difference scores were subjected to a non-parametric Friedman ANOVA. In case our analyses yielded significant effects, post-hoc tests were conducted. To determine whether potential differences in cardiovascular dynamics between UP and DOWN trials were associated with volitional pupil modulation performance at the beginning (i.e., day 1) and at the end of training (i.e., day 3), we calculated Spearman correlation coefficients between the pupil modulation index (i.e., UP–DOWN) and heart rate (UP–DOWN). Reported statistical analyses were two-tailed tests and corrected for multiple comparisons using sequential Bonferroni correction^76^.

### Experiment 1B: Replication of Experiment 1A

#### Participants

We recruited an independent cohort of 26 participants (18 females, 26 ± 7 years). Technical issues with the eye tracker led to an interruption of training and exclusion of one participant (n = 25 for final analyses).

#### Pupil-based biofeedback

The paradigm used was identical to Experiment 1A with two implemented changes: (i) we added a no feedback session *prior* to training (Pre) in which participants only used instructed mental strategies without receiving any feedback; (ii) online feedback on training day 3 was removed and participants only received post-trial performance feedback (SF 4A). As during no feedback trials, the baseline phase was indicated by a “=” above the fixation dot on the screen, changing to an “x” as soon as the modulation phase started. Trial timing and instructions remained the same as in Experiment 1A.

#### Cardiovascular measurements

For 15 participants (11 females, 25 ± 6 years), we additionally recorded cardiac and respiratory data with the same system and settings as described in Experiment 1A. Respiratory data is not reported here.

#### Pupil data (pre-)processing and analysis

The no feedback session (Pre and Post pupil-BF training) allowed us to directly compare self-modulation performance without feedback effects in the same group of participants. We excluded 5.4% of all trials due to violations during baseline and modulation phases (i.e., squinting/opening of the eye in a systematic way, large eye movements/saccades). Baseline-corrected pupil size time series during self-regulation were calculated as described in Experiment 1A. We statistically compared these time series Pre and Post pupil-BF training by subjecting the data to a two-way repeated measures ANOVA with the within-subjects factors *condition* (UP versus DOWN) and *session* (Pre versus Post pupil-BF training) using the MATLAB-based SPM1D toolbox for one-dimensional data (SPM1D M.0.4.8; https://spm1d.org/). SPM1D uses random field theory to make statistical inferences at the continuum level regarding sets of one-dimensional measurements. It is based on the idea to quantify the probability that smooth, random one-dimensional continua would produce a test statistic continuum whose maximum exceeds a particular test statistic value and has been previously used to analyze one-dimensional kinematic, biomechanical or force trajectories^81, 82^. In case of significant effects, post-hoc tests implemented in the SPM1D software were used which are corrected for multiple comparisons using Bonferroni corrections. In addition, we tested whether prior to pupil-BF training (Pre), pupil modulation index time series were already significantly different from 0 via a one-sample t-test against 0 implemented in the SPM1D toolbox. A significant result would indicate that participants were able to self-regulate pupil size to a certain extent already prior to training.

#### Cardiac data (pre-)processing and analyses

ECG R peaks were detected, extracted and further processed as described in Experiment 1A. Data of one participant was disrupted and excluded since it was not possible to detect R peaks for no feedback trials prior to training as well as on the first training session. Heart rate and RMSSD based on R-R-intervals were calculated. As and additional HRV measure, we further calculated the percentage of successive normal cardiac interbeat intervals greater than 35ms (pNN35) based on R-R intervals. The pNN35 is another common HRV measure and was used in other neurofeedback studies aimed at arousal regulation^67^. Heart rate and HRV (RMSSD and pNN35) calculated for each modulation phase was averaged across the respective up- and downregulation condition of each training session.

To investigate whether pupil self-regulation systematically modulates cardiovascular parameters, we subjected heart rate averages for each condition (UP versus DOWN) and training session (no feedback pre, day 1, day 2, day 3, no feedback post) to a two-way repeated measures ANOVA. Sphericity was assessed using Mauchly’s sphericity test and violations were accounted for with the Greenhouse-Geisser correction. Since our HRV measure, RMSSD, significantly deviated from normal distribution (Shapiro-Wilk tests, all *p <* 0.01), we calculated differences scores in RMSSD between up- and downregulation trials (DOWN-UP) for each session (no feedback pre, day 1, day 2, day 3, no feedback post). Positive values indicated larger HRV in DOWN than UP trials and negative values indicated larger HRV in UP than DOWN trials. These difference scores were subjected to a robust ANOVA using the WRS2 package of R. In case our analyses yielded significant effects, post-hoc tests were conducted. To determine whether potential differences in cardiac dynamics between UP and DOWN trials were associated with volitional pupil modulation performance prior to training, we calculated Spearman correlation coefficients between the pupil modulation index (i.e., UP–DOWN) and heart rate (UP–DOWN) or RMSSD (DOWN–UP). Reported statistical analyses were two-tailed tests and corrected for multiple comparisons using sequential Bonferroni correction^76^ or Hochberg’s approach implemented in the WRS2 package of R.

### Experiment 2: Pupil-BF combined with simultaneous fMRI

#### Participants

For Experiment 2, we re-recruited 25 participants (17 females; age 25 ± 5 years) from the pupil-BF groups of Experiments 1A and B. No statistical methods were used to pre-determine sample sizes but our sample size is similar or larger to those in previous fMRI studies showing significant correlations between pupil measures and brain activity^6, 8, 9^. Additional exclusion criteria were: (i) metal implants and/or fragments in the body, (ii) suffering from claustrophobia, and (iii) pregnancy at the time of the experiment. In case the last session of pupil-BF took place more than ten days before the scheduled fMRI session, participants received one session of re-training combining 40 trials of online feedback (i.e., 20 trials UP and DOWN, respectively) and 20 trials of only post-trial feedback (i.e., 10 trials UP and DOWN, respectively) in the week before the first fMRI session. This re-training was included to assure that participants were still able to modulate their own pupil size successfully. Re-training data is not reported here.

#### Data acquisition

All volunteers participated in two fMRI sessions, one using an optimized brainstem imaging sequence, the other acquiring whole-brain data (order of sessions was counterbalanced across volunteers). (f)MRI data was acquired on a 3T Philips Ingenia system using a 32-channel head coil. Anatomical images of the brain used for anatomical co-registration were acquired using a T1-weighted sequence (MPRAGE; 160 sagittal slices, TR/TE = 9.3/4.4 ms, voxel size = 0.7 mm^3^, matrix size = 240 × 240, flip angle = 8°, FOV = 240 mm (AP) × 240 mm (RL) × 160 mm (FH)). A turbo spin echo (TSE) neuromelanin-sensitive structural scan was acquired for delineation of the LC (not reported here) which was oriented perpendicular to the floor of the fourth ventricle (20 slices, 1.5mm, no gaps, in-plane resolution: 0.7 × 0.88 mm (reconstructed at: 0.35 × 0.35 mm), TR = 500 ms, TE = 10 ms, flip angle = 90°, covering the brainstem only). During the brainstem session, echo-planar imaging (EPI) scans were acquired in 39 sagittal slices (thickness: 1.8 mm, no gaps) aligned parallel to the floor of the fourth ventricle along the brainstem with the following parameters: TR = 2.5 s, TE = 26 ms, flip angle = 82°, SENSE acceleration factor = 2.1, in-plane resolution of 1.8 × 1.8 mm, and 240 volumes per run. Additionally, a whole-brain single volume EPI image was acquired to improve co-registration between fMRI data and the anatomical MRI scans (identical sequence parameters to the brainstem fMRI sequence).

During the whole-brain session, EPI images were acquired in 40 slices (thickness: 2.7 mm) with the following parameters: TR = 2.5 s, TE = 30 ms, flip angle = 85°, FOV = 223 mm (AP) × 223 mm (RL) × 116 mm (FH), Sense factor 2. Images were acquired at an in-plane resolution of 2.7 × 2.7 mm and were reconstructed at a resolution of 1.74 × 1.74 mm. In each session, we acquired 240 volumes per run. For the first seven subjects, we used an EPI sequence with slightly different details (i.e., especially higher spatial resolution to be able to see brainstem activity in small nuclei; non-isometric: 36 slices (thickness: 3mm, no gaps, TR = 2.5 s, TE = 30 ms, flip angle 82°, FOV = 210 mm (AP) × 210 mm (RL) × 108 mm (FH), Sense factor 2.2, in plane resolution of 2 × 2 mm reconstructed at a resolution of 1.64 × 1.64 mm), however, due to the strong distortion observed in one excluded participant, we decided to adjust the sequence.

During both MRI sessions, cardiac cycle and respiratory activity were recorded with a sampling rate of 496 Hz with a pulse oximeter attached to the left index finger and a chest belt, respectively (Invivo; Phillips Medical Systems, Orlando, Florida). These data were used for physiological noise removal as well as for calculating cardiovascular parameters.

Concurrently with fMRI recordings, participants’ eye gaze and pupil size of the left eye were tracked at 1000Hz using an EyeLink 1000 Plus Long Range Mount (SR Research, Ottawa, Ontario, Canada) that was placed at the end of the scanner bore on an aluminum stand and a first-surface mirror attached to the head coil. At the beginning of each fMRI run, the eye tracker was calibrated using 5-point calibration implemented in the software of the device. Pupil diameter was converted from arbitrary units to mm using the following formula: diameter (mm) = diameter (a.u.) / 1372. The conversion factor was estimated as described in ^70, 83^.

#### fMRI Paradigm

The pupil-BF paradigm during fMRI was similar to our training paradigm. Participants were instructed to maintain gaze on the green fixation dot presented in the center of the screen. Each trial began with a baseline measurement (7.5s) followed by an UP or DOWN phase (15s; without online feedback) during which participants were asked to apply the acquired respective mental strategy. Participants only received post-trial performance feedback (2.5s). After a short break (randomly jittered between 6-9s), the next trial started. Online processing and calculation of the feedback displayed to participants was performed as in Experiment 1A. The conditions were blocked within each of four fMRI runs, that is four trials of upregulation followed by four trials of DOWN (or vice versa). At the beginning of each block, it was indicated on the screen whether this block contains UP or DOWN trials. After each block, a break of 10s was added to allow participants to take short breaks in between. Each of the runs comprised two blocks of each condition, leading to 32 trials per condition. The order of conditions was counterbalanced across runs. Further, we counterbalanced across participants what conditions they started the respective fMRI session with.

#### Processing and analysis of pupil data

Pupil data (pre-)processing was conducted as described for Experiment 1A leading to baseline-corrected pupil diameter time series for UP and DOWN trials for each fMRI session. We statistically compared these time series for each session by subjecting the data to a paired-samples t-test (UP versus DOWN) using the SPM1D toolbox. Similar to Experiments 1AB, absolute baseline pupil diameter was compared by means of a repeated measures ANOVA with the within-subjects factor *session* (brainstem versus whole-brain) and *condition* (UP versus DOWN) and Bayesian repeated measures ANOVAs using default priors.

Finally, preprocessed absolute pupil diameter data were downsampled throughout the whole experiment to match the fMRI volume acquisition by averaging all data within a given TR. Missing pupil diameters were linearly interpolated across adjacent epochs resulting in a pupil diameter vector for each fMRI run.

#### Processing and analysis of cardiac data

Heart (pulse) rate concurrently measured during fMRI was (pre-)processed as described for Experiment 1B resulting in heart rate and HRV (RMSSD and pNN35) averages for each condition (UP and DOWN) and session (brainstem and whole-brain). Due to technical issues (whole-brain session of 1 participant), poor data quality (brainstem session of 1 participant), the final n for heart rate and HRV analyses was 24 for each fMRI session. Heart rate data was subjected to a repeated measures ANOVA with the within-subjects factor condition (UP versus DOWN) and fMRI session (brainstem vs. whole-brain). Since HRV values did significantly deviate from normal distribution (revealed by Shapiro-Wilk tests *p* < .05), we compared UP and DOWN trials for each fMRI session by means of a Wilcoxon signed rank test implemented in SPSS. Finally, as for the no feedback session prior to pupil-BF trraining, we calculated Spearman correlation coefficients between the pupil modulation index (UP– DOWN) and differences in heart rate (UP–DOWN) or RMSSD and pNN35 (DOWN–UP), respectively following training during fMRI (averaged across the two sessions). All statistical analyses were computed using SPSS and corrected for multiple comparison using sequential Bonferroni correction.

#### MRI analysis

MRI data was analyzed using tools from FSL version 6.0.5.2 (http://fsl.fmrib.ox.ac.uk/fsl), MATLAB R2018a, and FreeSurfer version 6.0 (https://surfer.nmr.mgh.harvard.edu/).

##### Pre-processing of brainstem data

Since brainstem fMRI signal is highly susceptible to corruption from physiological sources of noise, we used a specific pre-processing pipeline implementing suggestions from previously published research^6, 14, 84^ including the following steps: brain extraction using FSL’s automated brain extraction tool (BET;^85^), motion correction using the Linear Image Registration Tool (MCFLIRT;^86^), spatial smoothing using a 3mm full width-at-half-maximum (FWHM) Gaussian kernel, and a 90s high-pass temporal filter as implemented via FSL’s Expert Analysis Tool (FEAT;^87^). Additionally, each functional run showing excessive motion with an absolute mean displacement greater than 1 mm (i.e., ∼half a voxel size) was excluded (3 runs for 1 participant which led to exclusion from further analyses). To further de-noise the data, we performed Independent Component Analysis (ICA) and FLS’s Physiological Noise Modelling (PNM;^88^), an extended version of RETROICOR^89^). ICA denoising was performed using FSL’s Multivariate Exploratory Linear Optimized Decomposition Interface (MELODIC,^90^), which allows for decomposition of the fMRI data into spatial and temporal components. The resulting components were visually inspected and classified as “Noise” or “Signal” by two independent researchers using published guidelines^91^. Once labelled, the temporal information of the noise components was extracted and ICA noise regressor lists were generated for later implementation in our general linear model (GLM). For PNM, cardiac and respiratory phases were assigned to each volume in the concatenated EPI image time series separately for each slice. Our complete physiological noise regression model included 34 regressors: 4^th^ order harmonics to capture the cardiac cycle, 4^th^ order harmonics to capture the respiratory cycle, 2^nd^ order harmonics to capture the interaction between cardiac and respiratory cycles, 2^nd^ order harmonics to capture the interaction between respiratory and cardiac cycles, one regressor to capture heart rate, and one regressor to capture respiration volume. Subsequently, we visually inspected the waveforms to assure that the peak of each respiratory and cardiac cycle was correctly marked and adjusted if necessary. The resulting voxelwise confound lists were later added to our GLM. Note that physiological data acquisition was corrupted for one participant. Therefore, this data was analyzed without the PNM regressors. Image co-registration from functional to Montreal Neurological Institute (MNI) standard space was performed by (i) aligning the functional images of each run to each subject’s whole-brain EPI image, using the Linear Image Registration Tool (FLIRT) employing a mutual information cost function and six degrees of freedom^92^; (ii) registering whole-brain EPIs to structural T1-weighted images, using a mutual information cost function and six degrees of freedom, and then optimised using boundary-based registration (BBR;^93^); (iii) co-registering structural images to the MNI152 (2mm^3^) template via the nonlinear registration tool (FNIRT) using 12 degrees of freedom^94^, and applying the resulting warp fields to the functional images. Each co-registration step was visually inspected using Freeview (FreeSurfer) to ensure exact alignment with an emphasis placed on the pons regions surrounding the LC. Unfortunately, 3 out of 25 participants needed to be excluded from brainstem fMRI analyses due to heavy distortions (including brainstem regions; 1 participant), excessive motion (1 participant, see above), and periods of sleep during the measurement (1 participant). As additional control analyses to exclude smearing of noise from the 4^th^ ventricle to the brainstem, we abstained from spatially smoothing the data during pre-processing, while all other (pre)processing steps remained the same (for results, see SF 6).

##### Pre-processing of whole-brain data

We used a similar pre-processing pipeline as described above, however, with the following differences: Spatial smoothing was applied using a 5mm full width-at-half-maximum (FWHM) Gaussian kernel. Functional EPI-images were aligned to structural T1-weighted images using linear registration (FLIRT). Structural images were aligned to the MNI standard space using nonlinear registration (FNIRT), and the resulting warp fields applied to the functional images. Data from 1 out of 25 participants showed very strong distortions in temporal and frontal regions and was excluded from further analysis. Also, we needed to exclude 1 functional run for 3 participants and 2 functional runs for 1 participant due to excessive motion with an absolute mean displacement greater than half a voxel size (i.e., 1.4 mm). We did not apply PNM to the data and, as recommended by FSL (https://fsl.fmrib.ox.ac.uk/fsl/fslwiki/FEAT/UserGuide), ICA was only performed (n = 9) if residual effects of motion were still left in the data even after using MCFLIRT for motion correction.

##### fMRI analyses

Time-series statistical analysis were performed using FEAT with FMRIB’s Improved Linear Model (FILM). Two equivalent GLMs were estimated for each session (i.e., for the whole-brain session and the brainstem session).

First, we directly contrasted brain activity during up-versus downregulation periods during whole-brain scans. Therefore, “UP” and “DOWN”, were modelled as main regressors of interest for the complete modulation phase of 15s. Instruction, baseline, and post-trial performance feedback periods were modelled as separate regressors. All regressors were convolved with a double gamma HRF and its first temporal derivative. White matter (WM) and cerebrospinal fluid (CSF) time-series, motion parameters and ICA noise component time-series (if applied), respectively, were added as nuisance regressors. Second, we investigated which brain areas would change their activity in close association with pupil diameter. Therefore, pupil diameter was entered as a regressor of interest, together with the same nuisance regressors as described above. To adjust for delays between brain and pupillary responses, we shifted the pupillary signal by 1s.

For the brainstem session, we defined GLMs with identical regressors that were, however, convolved with the default optimal basis sets (i.e., the canonical hemodynamic response function (HRF) and its temporal and dispersion derivatives) using FMRIB’s Linear Optimal Basis Sets toolkit^95^. This convolution was chosen to account for the possibility that brainstem BOLD responses such as in the LC are not well-modelled solely by the canonical HRF^96^. However, to avoid overestimating statistical effects, only the canonical HRF regression parameter estimates were used for subsequent higher-level analyses. Additionally, the PNM voxelwise confound lists and the time series of ICA noise components lists (but not motion parameters) were used as nuisance regressors.

All described GLMs were computed separately for each run of each participant and then averaged across runs for each participant in second level analyses using FMRIB’s Local Analysis of Fixed Effects^97^. To determine significant effects at the group level, a third level mixed-effects analyses was performed. Group z-statistic images were thresholded at z > 3.1 for whole-brain analyses and family-wise-error corrected (FWE) at the cluster level. For the brainstem analysis we report results thresholded at z > 2.3, FWE-corrected at the cluster level. To visualize brainstem activity, we created a mask covering the brainstem (derived from the Harvard-Oxford subcortical atlas; thresholded at 0.5 and binarized using FSL’s image calculator fslmaths) and the NBM (Ch4 cell group derived from the JuBrain/SPM Anatomy Toolbox^98^), showing z-statistic images within these predefined regions.

##### Regions of Interest (ROI) analysis

We conducted a series of ROI analyses, to test our hypothesis that self-regulation of pupil size is linked to activity changes in the LC and other neuromodulatory, arousal-regulating centres in the brain. Our principal region of interest was (i) the LC, however, given the close interaction with other brainstem areas and neuromodulatory systems involved in pupil control, we defined secondary ROIs, namely (ii) the cholinergic NBM, (ii) the dopaminergic SN/VTA, (iii) the serotonergic DRN, and (iv) the SC – a specialized motor nucleus in the midbrain implicated in the pupil orientation response – as a control region. Probabilistic anatomical atlases were used to define the location of the brainstem ROIs^43–49^ and the NBM (Ch4 cell group^98^; see Fig. 3E) in MNI space. After careful visual inspection, we decided to threshold the SC and DRN mask at 0.1, and the SN/VTA-mask at 0.3. The masks for all ROIs were binarized. In a next step, we extracted the average mean signal intensity (z-values) within each ROI from the second-level fixed effects analysis conducted for each participant for each GLM (i.e., for UP versus rest and DOWN versus rest, and pupil diameter vector used as regressor). Due to deviation from normal distribution for the NBM (Up>Rest; Shapiro-Wilk tests; *p <* .05) and SC (pupil regressor; Shapiro-Wilk tests; *p <* .05), we used Wilcoxon signed rank tests to test for a significant difference between z-values for UP and DOWN periods within the NBM, as well as for a significant correlation between absolute pupil diameter and BOLD activity within the SC (i.e., deviating from 0). For all other ROIs, we used paired samples t-tests and one sample t-tests against 0. All analyses were corrected for multiple comparisons using sequential Bonferroni correction. Even though we conducted pre-processing using PNM and ICA, we performed additional control and specificity analyses. To this end, we used additional control masks covering (i) the whole brainstem including the midbrain, pons, and the medulla oblongata, and (ii) the 4^th^ ventricle (SF 5A). First, as for our ROIs, we extracted the average mean signal intensity (z-values) within each ROI from the second-level fixed effects analysis conducted for each participant for each GLM (i.e., for UP versus rest and DOWN versus rest) and used two-tailed Wilcoxon signed tank tests to test for a significant difference between z-values for UP and DOWN periods in these additional control regions. Further, all ROI analyses were repeated for data without spatial smoothing applied during preprocessing.

### Experiment 3: Pupil-BF combined with an auditory oddball task

#### Participants

Experiment 3 involved 22 participants (15 females; 26 ± 6 years) of the 25 participants re-recruited for the fMRI experiment. Unfortunately, 3 participants dropped out because of personal reasons. Participants received one session of re-training combining 40 trials of online feedback (i.e., 20 trials UP and DOWN, respectively) and 20 trials of post-trial feedback (i.e., 10 trials UP and 10 DOWN, respectively) within 7 days prior to Experiment 3, in case the last session of pupil-BF took place more than ten days before the scheduled pupil-BF oddball session. This re-training was included to assure that participants were able to modulate their own pupil size successfully. Re-training data is not reported here.

#### Paradigm: Pupil-BF with an integrated auditory oddball task

The auditory oddball task during pupil-BF consisted of 186 target (∼20%) and 750 standard (∼80%) tones (440 and 660 Hz with the pitch for target being counterbalanced across participants) delivered for 100ms via headphones at approximately 60dB (Panasonic RP-HJE125, Panasonic Marketing Europe GmbH, Wiesbaden, Germany), and was embedded in our pupil-BF paradigm during which participants were volitionally up- or downregulating their own pupil size via acquired mental strategies. In addition to these two self-regulation conditions, we added a third control condition in which participants were asked to count back in steps of seven instead of self-regulating pupil size. To ensure that participants stick to this task, they were occasionally prompted to enter the final number (after every 6^th^ counting trial). This condition was added to control for effort-effects related to pupil self-regulation and to check whether oddball responses differ in self-regulated as compared to more natural fluctuations in pupil size. At the beginning of the task, participants were reminded to focus on their mental strategies and to pay attention to the tones at the same time.

Each pupil-BF-oddball trial began with a short instruction (2s) indicating the required pupil self-regulation direction (i.e., UP or DOWN) or the counting control condition followed by a baseline measurement of 4s in which participants were asked to silently count backwards in steps of seven. Then, participants were asked to apply the acquired mental strategy to up- or downregulate pupil size for 18s or to continue counting (i.e., control). During this self-regulation or counting phase, eight tones (one or two targets and seven or six standards) were played, and participants were asked to respond to targets but not to standards by pressing a key as quickly as possible with the index finger of their preferred hand. The first tone which was always a standard in each self-regulation or counting phase was played after the first jitter phase (1.8 – 2.2s after the baseline phase) to ensure that participants already reached some modulation of pupil size before the first target appeared. The inter-stimulus interval (ISI) between tones was randomly jittered between 1.8 – 2.2s. Between two targets, there were at least two standards presented leading to a minimum ISI of 5.4s between two targets allowing the target-evoked pupil dilation response to return to baseline between targets. After each trial, participants received post-trial feedback on average and maximum pupil diameter change for 2s. After a short break of 2s, the next trial started. The calculation of feedback was similar to Experiment 1A, except that only the last two seconds of the baseline phase were considered due to the shortened baseline phase of 4s. For counting trials without feedback phases, counting was followed by a 4s break to keep trial timings similar. Instructions and indication of task phases on the screen were displayed as in Experiment 1B on Day 3. However, to improve visibility further, indication of task phase changes from baseline measurements to self-regulation was displayed in magenta (instead of green) isoluminant to the gray background (Fig. 6A).

The pupil-BF conditions were blocked, each block containing 13 trials of UP, DOWN, and control, respectively. The number and order of tones was the same for each pupil-BF condition. After each block, participants could take a self-paced break. In total, the task consisted of nine blocks leading to 117 pupil-BF trials in total. The order of blocks was pseudorandomized in the sense that each condition needed to be presented once (or twice) before another condition was presented for the 2^nd^ (or 3^rd^) time.

#### Physiological data acquisition

Eye gaze, pupil diameter, cardiac and respiratory data were recorded throughout the task as described in Experiment 1A and B. Additionally, we acquired electroencephalography (EEG) data using 64 Ag/AgCl actiCAP active surface electrodes, an actiCHamp Plus amplifier and the Brain Vision Recorder proprietary software (Brain Products, Munich, Germany). However, cardiac, respiratory and EEG data are not reported here.

#### Data (pre-)processing and statistical analyses

##### Pupil-BF: Self-regulation of pupil size under dual task conditions

Pupil data (pre)processing was conducted as described for Experiment 1A. We had to exclude 2 participants from further analyses leading to a final n = 20 for pupil data analyses: one participant showed a very high blink rate during the oddball with more than 30% of missing samples during baseline and self-regulation or counting phases for more than 50% of the trials. A second participant showed a similar pattern with more than 30% of missing samples in 35% of the trials, mainly in the control condition (only 20% of the trials were usable after pre-processing). Combined with low behavioral performance (correct responses to targets deviated more than 3 SD from the group mean), we decided to exclude this participant from pupil and behavioral data analyses. For each of the remaining participants, we derived *absolute* and *baseline-corrected* pupil diameter time series averaged for UP, DOWN or counting trials throughout the baseline and self-regulation or counting phases of the pupil-BF oddball task. Both absolute and baseline-corrected pupil diameter time series were extracted since we expected to observe target-evoked pupil dilation responses that depend in size on absolute pupil size prior to tone onset. Baseline correction was applied as described above, choosing a baseline of 1s prior to self-regulation and counting, respectively.

For statistical analyses of self-regulated pupil size, we averaged preprocessed, baseline-corrected pupil diameter time series for each condition across the entire self-regulation or counting phase to test whether participants were able to successfully self-regulate pupil size even under challenging dual-task conditions. Since these data were deviating from normal distribution (Shapiro Wilk test; *p* < 0.05), we subjected the data to a robust repeated measures ANOVA implementing 20% trimmed means with the factor *condition* (UP versus DOWN versus control; R package WRS2). In case of a significant main effect, we computed the corresponding post-hoc tests included in this R package with implemented FWE correction. Next, we averaged absolute pupil diameter samples across the last second of the baseline phase and subjected it to a repeated measures ANOVA with the within-subjects factor *condition* (UP versus DOWN versus control) to compare absolute baseline pupil diameter between the three conditions. Sphericity was assessed using Mauchly’s sphericity test and violations were accounted for with the Greenhouse-Geisser correction. In case of a significant effect, post-hoc tests were conducted using the sequential Bonferroni correction for multiple comparisons. In case of no significant effect, we ran an analogous Bayesian repeated measures ANOVA using JASP (with default priors) to evaluate whether there is evidence for the null hypothesis (i.e., no baseline differences between conditions). All reported statistical analyses were two-tailed tests.

##### Pupillary responses to presented tones during the oddball task: Pre-tone baseline pupil diameter and pupil dilation responses

We extracted 3.5s epochs around each standard or target tone from -0.5 to +3s relative to tone onset. Pre-tone pupil diameter was calculated by averaging 500ms of pupil diameter data preceding tone presentation. The pupil time series was then (i) normalized to this pre-tone pupil diameter baseline and (ii) averaged for each stimulus type (standard versus target) and condition (UP versus DOWN versus control).

To compare *target*-evoked pupil dilation responses between conditions, we subjected the baseline-corrected pupil dilation response time series to targets grand-averaged within each condition to a repeated-measures ANOVA with the within-subjects factor *condition* (UP versus DOWN versus control) using the SPM1D toolbox. This time series data analysis allowed us to compare both peak dilation towards targets and the shape of the dilation response statistically. The same analysis was repeated for pupil dilation responses to standard tones as a control analysis. In case of significant main effects, post-hoc tests implemented in the SPM1D software corrected for multiple comparisons using Bonferroni correction were used.

##### Behavioral data

Throughout the experiment, we recorded the correctness of responses and reaction times to tones. Responses to standard tones (i.e., false alarms) and responses for which reaction times exceeded two SD below or above the mean of the respective participant were excluded from further reaction time analyses. Lastly, we calculated the percentage of hits (i.e., responses to targets) and false alarms (i.e., responses to standards) and obtained a percentage of correct responses (i.e., accuracy) by subtracting these two measures from each other^64^.

To investigate whether self-regulation of pupil size modulates behavioral task performance on the oddball task, we subjected individual reaction times to target tones averaged across trials of each condition to a repeated measures ANOVA with the within-subjects factor *condition* (UP versus DOWN versus control). Sphericity was assessed using Mauchly’s sphericity test and violations were accounted for with the Greenhouse-Geisser correction. Next, we examined whether not only speed per se but also performance variability indicated by the standard deviation (SD) of reaction times may be modulated by self-regulated pupil size. We subjected the SD of reaction times to a repeated measures ANOVA with the within-subjects factor *condition* (UP versus DOWN versus control). Sphericity was assessed using Mauchly’s sphericity test and violations were accounted for with the Greenhouse-Geisser correction. In case of a significant effect, we conducted post-hoc tests controlled for multiple comparisons by means of sequential Bonferroni corrections. Reported statistical analyses were two-tailed tests.

##### Link between pupil and behavioral data

In a next step, we computed repeated measures correlation coefficients between the derived single-trial pre-tone absolute baseline pupil diameter (averaged 500ms before target onset) and baseline-corrected target-evoked pupil dilation response peaks of all participants using the R package rmcorr^99^. Since the correlation between a variable × (i.e., baseline pupil diameter) and the variable y-x (i.e., baseline-corrected pupil dilation response peaks) is susceptible to a regression towards the mean effect, we decided to use relative instead of subtractive baseline correction for these additional analyses (i.e., 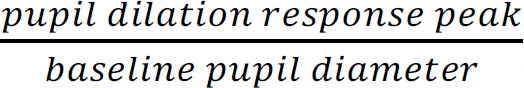). Even though correlating a relative change from a variable to this variable itself is also not free of bias, it is indeed possible to test for an unbiased relationship by fitting the following model in R for y = Pupil Dilation Response Peak and x = Baseline Pupil Size and consider the resulting coefficient

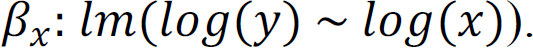

This model corresponds to the following equation:

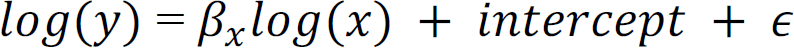

which is equivalent to:

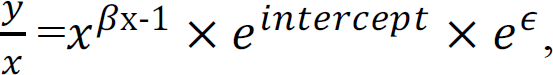

Importantly, this is an unbiased estimate of an influence of the baseline pupil size x on the following pupil dilation response peak y/x (i.e., randomly generated estimates of β by simulations center around 0). Then, we tested with a one-sample *t-*test the H_0_: β_1_ − 1 = 0, i.e., that baseline pupil size has no significant influence on pupil dilation responses.

Finally, to investigate the relationship between pupil size measures and behavioral responses at a single trial level, we conducted additional repeated measures correlation between reaction times and baseline pupil size and pupil dilation responses, respectively. Since both baseline pupil diameter and reaction times to targets could potentially be influenced by previously reported time-on-task effects^61, 100^, we ran linear mixed models on single-trial level similar to Arnau et al.^101^ with the dependent variables reaction times and baseline pupil size, respectively. Time-on-task entered the model as a fixed effect and was operationalized as the number of the respective trial in the experiment in ascending order. A random intercept was modeled for each participant (i.e., reaction time ∼ time on task + (1|participant) and baseline ∼ time on task + (1 |participant), respectively). In case of significant time-on-task effects, we removed a linear trend in the data for each participant using the *detrend* function in MATLAB (method: linear) before subjecting it to repeated measures correlations in R.

## Data availability

Processed pupil, fMRI, heart rate, heart rate variability, and behavioral data will be made openly available on the ETH Library Research Collection (including a DOI).

## Code availability

Unfortunately, scripts used for execution of the pupil-based biofeedback cannot be made publicly available. The exact code of the pupil-based biofeedback algorithm is proprietary software of ETH Zurich and cannot be shared beyond the description of the algorithm given in the methods section.

## Acknowledgments

The authors would like to express their gratefulness to all participants of the study. We further thank J.W. de Gee for his great help regarding brainstem-specific MRI sequences, V. Zerbi, J. Bohacek, J.M. Shine and O. Harrison for fruitful discussions, Lukas Graz for his statistical advice, and W. Potok and D. Woolley for their valuable help and feedback on the manuscript. S.N.M. discloses support for the research of this work from the Swiss National Science Foundation (SNSF) Spark grant (CRSK-1_190836) and is supported by the Hochschulmedizin Zürich Flagship project STRESS. J.I. is supported by SNSF grant 32003B_207719. N.W. is supported by the National Research Foundation, Prime Minister’s Office, Singapore under its Campus for Research Excellence and Technological Enterprise (CREATE) programme and funded by the SNSF and Innosuisse BRIDGE Discovery grant (40B2-0_203606). The funders had no role in study design, data collection and analysis, decision to publish or preparation of the manuscript.

## Author Contributions

S.N.M, M.B., and N.W. were involved in conceptualization and design of the study, S.N.M., J.I., and S.M., acquired the data, S.N.M., S.K., and N.W. planned the analysis, S.N.M, J.I., S.M, and M.C. analyzed the data; S.N.M and N.W. interpreted the data, S.N.M drafted the manuscript and M.B., S.K., J.I., S.M., M.C. and N.W. substantively revised it.

## Competing Interests

The authors declare the following competing interests: S.N.M, M.B, and N.W. are founders and shareholders of a company, MindMetrix, and have a patent application related to used methods (patent applicant: ETH Zurich; inventors: M.B, S.N.M, N.W., pending patent applications EP21704565.7 and US17/800,455). All other authors declare no competing interests.

## Notes

### Competing Interest Statement

S.N.M, M.B, and N.W. are co-founders of a company (MindMetrix) that intends to offer related technology to the consumer market and have a patent application related to used methods pending (patent applicant: ETH Zurich; inventors: M.B, S.N.M, N.W.). All other authors declare no competing interests.

### Summary of Updates

Conducted additional control analyses and an additional control experiment (mainly in Figures in Supplementary File); corrected an error in Figure 2; changes in analyses of data displayed in Figure 6F

