## Supplementary Information for "Self-regulating arousal via pupil-based biofeedback"

### Supplementary Material

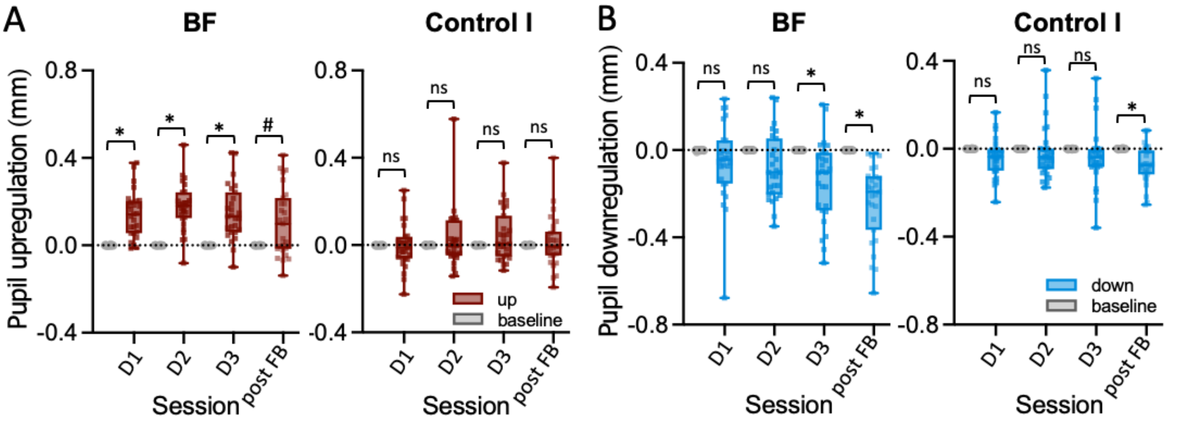

**Supplementary Figure 1.** Baseline-corrected pupil up- (A) and downregulation (B) averaged across the 15s modulation phase for each condition as compared to baseline pupil size in the last second before modulation for the pupil-BF and control group I, respectively. Whereas in the pupil-BF group upregulation was significantly different from baseline from D1 onwards, downregulation was significantly different only on D3 and during no feedback sessions after training (Wilcoxon signed-rank tests UP D1: z = -4.36; $p$ = .014; r = -0.84; D2: z = -4.42; $p$ = .015; r = -0.85; D3: z = -4.23; $p$ = .013; r = -0.81; no feedback: -2.64; $p$ = 0.08; r = -0.47; DOWN D1: z = 1.73; r = 0.33; D2 = 2.40; r = 0.46; all $p$ > 0.12; D3: z = 3.39; $p$ = .011; r = 0.65; no feedback: z = 4.54; $p$ = .016 r = 0.87). In the control group, upregulation was not significantly different on any of the training/no feedback sessions (UP D1: z = 0.65; r = 0.13; D2: z = 0; r = 0; D3: z = -1.25; r = -0.24; no feedback: z = -0.24; r = -0.05; all $p$ > 0.51). For downregulation, there was an only a significant effect for no feedback trials after training, even though all self-regulation values were lower in magnitude than in the pupil-BF group (DOWN D1: z = 2.52; r = 0.49; D2: z = 1.61; r = 0.31; D3: z = 2.40; r = 0.46; all $p$ > 0.10; no feedback: z = 3.41; $p$ = 0.012; r = 0.65). *$p$ < 0.05; # $p$ < 0.1. All p-values are corrected for multiple comparisons using the sequential Bonferroni procedure.

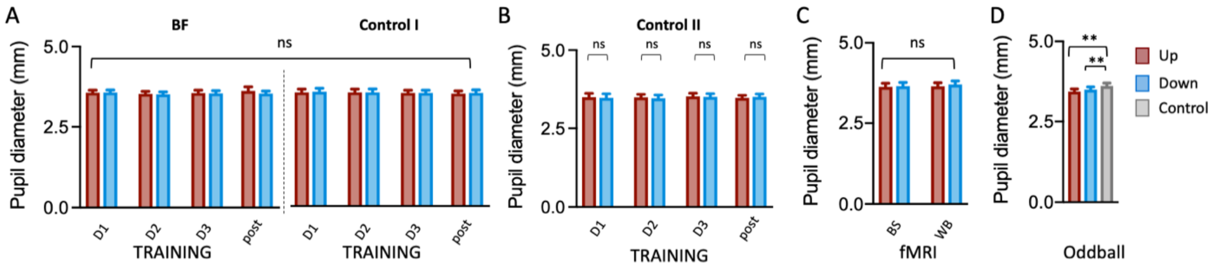

**Supplementary Figure 2.** Absolute baseline pupil diameter averaged across the last second of the baseline phase before modulation onset for pupil-BF training **(A and B)**, brainstem (BS) and whole-brain (WB) fMRI **(C)**, and the oddball task sessions **(D)**. Only during the oddball task, baseline pupil sizes differed between conditions (repeated measures ANOVA: $F(2,38) = 11.41$; $p < .001$) which was mainly driven by significantly larger pupil sizes in control than up- ($t(19) = -3.95$; $p = .003$) and downregulation trials ($t(19) = -3.43$; $p = .006$; sequential Bonferroni-corrected for multiple comparisons). All other analyses (i.e., pupil-BF training and fMRI sessions) did not reach significance regarding the factor *condition* (pupil-BF v Control I: all $p \geq .48$; Control II: all $p \geq .33$; fMRI: all $p \geq .09$). These findings were supported by additional Bayesian analyses providing anecdotal to moderate evidence in favor of H0, i.e., no significant difference between conditions for pupil-BF training (**A.** factor condition (up versus down): BF10 = 0.11; with an error percentage of 2.27%; similarly, there was no evidence for an interaction between conditions and training sessions: BF10 = 0.001; error percentage of 2.37%; **B.** up versus down on day 1: BF10 = 0.29; error% = 0.013; day 2: BF10 = 0.39; error% = 0.017; day 3: BF10 = 0.27; error% = 0.012 and no feedback post: BF10 = 0.29; error% = 0.013), and fMRI (**C.** factor condition (up versus down): BF10 = 0.28; error percentage = 1.15%; conditions * fMRI sessions: BF10 = 0.02; error percentage = 2.04 %).

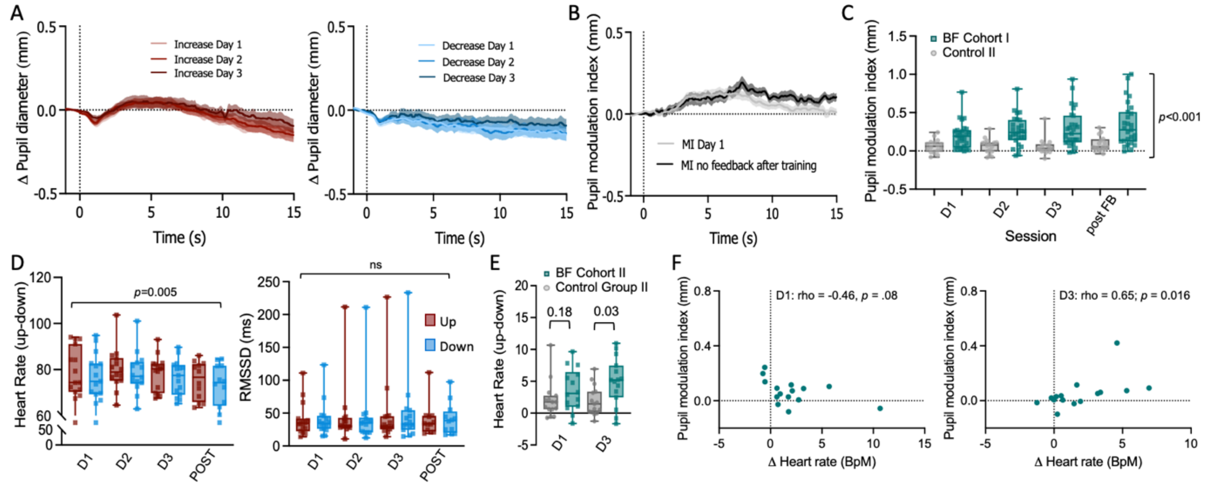

**Supplementary Figure 3.** Replication of Pupil-BF training results and effects on cardiovascular parameters in an independent 2nd cohort of control participants (Controls II; n = 16, 13 females, 25.06 ± 7.38 years) who received yoked biofeedback but believed it was veridical. **A.** Average changes in pupil size during 15s upregulation (UP) (left panel) and downregulation (DOWN) (right panel) are shown for Controls II for training sessions on Day 1, Day 2 and Day 3 of Experiment 1A. Please note that, descriptively, participants of control group II show some ability to self-regulate pupil size even when receiving yoked feedback. However, like in our initial control group (as depicted in Figure 2), this ability is reduced as compared to our pupil-BF group **B.** Time series of pupil modulation index reflecting the difference between up- and downregulation time series measured during the first day of training (day 1) and at the end of training during the no feedback session of Controls II. **C.** The pupil modulation index reflects the difference between the average pupil size during the two conditions (UP–DOWN) and is shown for each session (Day 1, Day 2, Day 3, no feedback post training session) and group (pupil-BF group of Experiment 1A and control group II; dots and squares represent individual participants). Pupil modulation indices were generally higher in the pupil-BF group compared to control group II replicating our results of control group I (robust ANOVA, main effect of group: $F(1,20.26) = 19.80$; $p < 0.001$; $\eta_p^2 = 0.49$; 95%-CI$\eta_p^2$ [0.16; 0.67]). The group*condition interaction was not significant ($p = 0.21$). **D** Heart rate (left panel) and heart rate variability (HRV) data (right panel) averaged for UP and DOWN trials across all participants of control group II during pupil-BF training (day 1: n = 15, day 2: n = 14; day 3: n = 15; no feedback post training: n = 12). HRV was estimated as the root mean square of successive differences (RMSSD). We found significantly higher heart rate during UP as compared to DOWN trials (significant main effect, repeated measures ANOVA: $F(1,10) = 12.88$; $p = 0.005$; $\eta_p^2 = 0.56$; 95%-CI$\eta_p^2$ [0.08;0.75]), however, this difference did not significantly change with training (i.e., no significant *condition*session* interaction ($p = 0.87$); no significant main effect of session: $p = 0.36$). Pupil self-regulation training did not significantly modulate HRV (Friedman ANOVA on DOWN-UP differences: $p = 0.98$). Further, testing for differences in HRV between DOWN and UP trials within each session did not yield significant results (Wilcoxon Signed-Rank tests: day 1: $p = 0.39$; day 2: $p = 0.36$; day 3: $p = 0.36$; no feedback: $p = 0.70$). **E.** Results of additional analyses comparing heart rate difference scores (UP-DOWN) between the control group II and the pupil-BF group at the beginning of (i.e., day 1) and at the end of training (day 3) via independent samples t-tests. Difference scores did not differ between groups on day 1 ($t(27) = -1.38$; $p = 0.18$; $d = -0.51$; 95%-CI$d$ [-1.25;0.23]). On day 3, this difference was, however, significantly larger in the pupil-BF as compared to the control group ($t(28) = -2.63$; $p = 0.03$; $d = -.096$; 95%-CI$d$ [-1.71;-0.19]; corrected for multiple comparisons) potentially indicating a learning effect in our cardiovascular measurement in the pupil-BF as compared to our control group**.** **F.** Spearman rho correlation coefficients between pupil modulation indices (i.e., the difference between pupil diameter changes in the two conditions; UP-DOWN) and differences in heart rate in Controls II (UP-DOWN) revealing a significant link between better pupil self-regulation and larger heart rate differences at the end (right panel; rho = 0.65; $p = 0.016$; 95%-CIrho [0.20;0.88]) but not at the beginning of training (left panel; only trend-level for negative correlation; rho = -0.46; $p = 0.08$; 95%-CIrho [-0.80; 0.08]). Error bars indicate SEM. All reported $p$-values are corrected for multiple comparisons. Shaded areas and error bars indicate SEM.

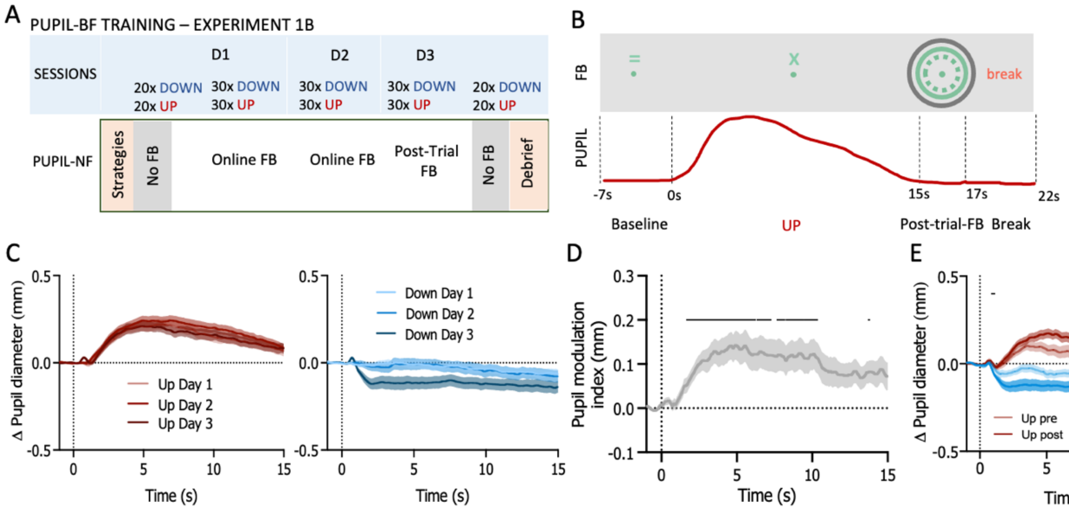

**Supplementary Figure 4.** Replication of Pupil-BF training results with an independent 2nd cohort of participants in Experiment 1B. **A.** As in Experiment 1A, healthy volunteers were informed about potential mental strategies of arousal regulation and then participated in 3 days (D1, D2, D3) of up-regulation (UP) and downregulation (DOWN) training (30 trails each) while receiving pupil-BF. Both before (Pre) and after (Post) training, all participants performed 20 UP and DOWN trials without receiving any feedback. At the end of training, all participants indicated used strategies during debriefing. **B.** Trial design and used colors were similar as in Experiment 1A, however, during the 15s modulation phase on Day 3, participants did not receive online feedback but only 2s of color-coded post-trial performance feedback (green = average circle size during modulation; black = maximum (UP) or minimum (DOWN) circle size during modulation). The change from baseline to modulation phases was indicated by a green '=' above the fixation dot changing to an 'x'. **C.** Average changes in pupil size during 15s upregulation (UP) (left panel) and downregulation (DOWN) (right panel) are shown for the 2nd cohort of the pupil-BF group (n = 25) for the training sessions on Day 1, Day 2, Day 3 of Experiment 1B **D.** Time series of the pupil modulation index measured during the no feedback session before pupil-BF training in Experiment 1B (independent cohort, n = 25). Solid black lines at the top indicate clusters of significantly higher modulation indices than 0 already prior to pupil-BF training (SPM1D one sample *t*-test; all $p < 0.05$). **E.** Pupil self-regulation time series for up- and downregulation trials pre and post training (i.e., no feedback trials). A repeated measures ANOVA with the factors *condition* and *session* implemented in SPM1D revealed a significant main effect of *condition* driven by stronger down- as compared to upregulation for only two short time windows during pupil self-regulation: at the beginning (0.93-1.05s; $p = .047$) and the very end of every modulation phase (14.8-15s; $p = .048$, significant clusters are indicated by the black solid lines above the time series; significant main effect of *session* is reported in the main results). However, this effect was independent of the respective session (no significant interaction) and may thus rather represent an inherent feature of pupil modulation with a slower build-up and a faster decline for up- as compared to downregulation trials. Shaded areas and error bars indicate SEM.

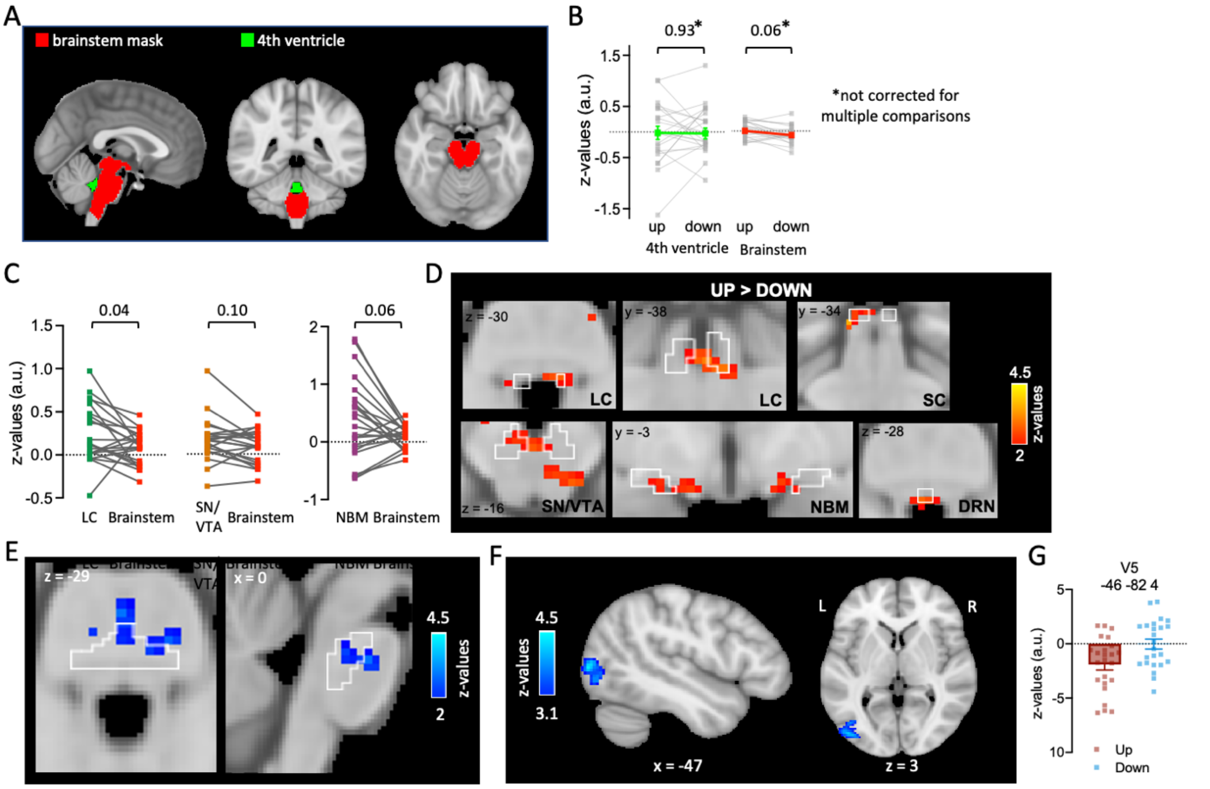

**Supplementary Figure 5.** Results of additional fMRI analyses on control regions in the brainstem and 4th ventricle as well as UP>DOWN and DOWN>UP results for brainstem and whole-brain sessions. **A.** Masks of additional control regions covering the complete brainstem (red) and the 4th ventricle (green). **B.** Activity during UP versus DOWN phases of pupil self-regulation extracted from the control regions as depicted in A. Statistical comparisons using paired *t*-tests revealed no significant effects of pupil self-regulation (UP>rest versus DOWN>rest) on BOLD activity extracted from each of the masks at an uncorrected level (only trend-level for complete brainstem: $t(21) = 2.00$; $p = 0.06$; $d = 0.43$; 95%-CId [-0.02; 0.86]; 4th ventricle: $t(21) = 0.09$; $p = 0.93$; $d = 0.02$; 95%-CId [-0.40; 0.44]; all *p*-values uncorrected). For the 4th ventricle mask, we even found moderate evidence for H0 when conducting additional Bayesian post-hoc analyses (4th ventricle: BF10 = 0.224; moderate evidence for H0, error % = 0.02; complete brainstem: BF10 = 1.18; anecdotal evidence for H1; similar to additional post-hoc Bayesian analyses that revealed moderate evidence for H0 for our initial ROIs SC: BF10 = 0.34 and DRN: BF10 = 0.31 as depicted in Figure 3B in the main manuscript). **C.** To investigate post-hoc whether the significant effects observed in our pre-defined ROIs (i.e., in the LC and SN/VTA; no significant but trend-level effects in the NBM) exceed those in the complete brainstem, we compared difference scores (UP>rest minus DOWN>rest: UP-DOWN) between these regions. Wilcoxon Signed Rank tests comparing the UP-DOWN difference revealed that only in the LC but not in the SN/VTA or NBM (only trend-level), the difference in BOLD activity between UP and DOWN is significantly larger than in the complete brainstem (LC: $z = 2.45$; $p = 0.04$; r = 0.52; SN/VTA: $z = 1.93$; p = 0.10; r = 0.41; NBM: $z = 1.87$; $p = 0.06$; r = 0.40; Bonferroni-Holm corrected). **D.** Data-driven analysis across the brainstem revealed a significant bilateral activation cluster in the LC, SN, VTA, NBM, and small parts of the SC and DRN during pupil upregulation (UP > DOWN), however, only at an uncorrected level ($p < 0.05$). For other regions involved in arousal and autonomic regulation activated by pupil up- as compared to downregulation, see ST 2A. Results for downregulation (DOWN > UP) in the brainstem **(E)** and whole-brain session **(F, G).** In the brainstem, we only found activation clusters in white matter tracts of the brainstem (for instance, the pontine crossing tract) at an uncorrected level ($p < .05$). During the whole-brain session, we found significantly higher BOLD activity during DOWN than UP in V5. **G.** Estimated BOLD response represented by z-values for UP versus rest and DOWN versus rest extracted from the peak voxel of the significant cluster shown in B. Please note that whole-brain activation maps are thresholded at $z > 3.1$ and FWE-corrected using a cluster significance level of $p < .05$. Locus coeruleus (LC); substantia nigra (SN) and ventral tegmental area (VTA) combined into one ROI (SN/VTA); dorsal raphe nucleus (DRN); nucleus basalis of Meynert (NBM); and superior colliculus (SC).

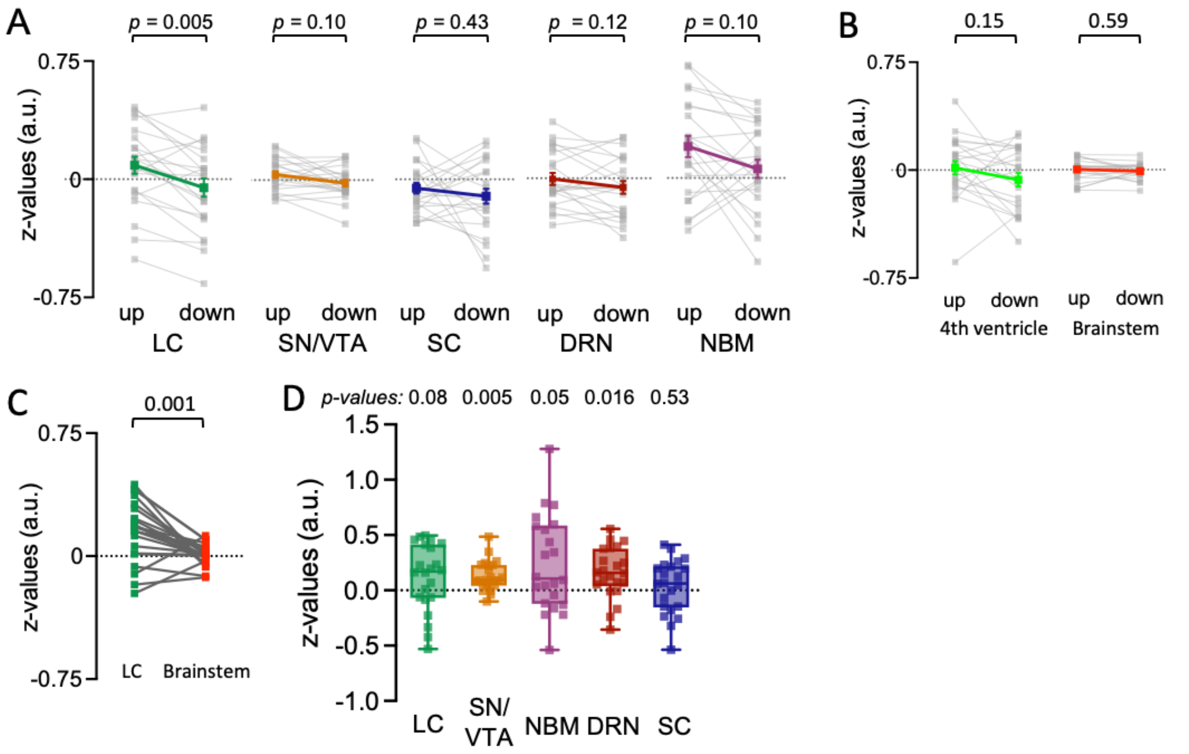

**Supplementary Figure 6.** Brainstem fMRI results of data without the application of spatial smoothing during preprocessing, **A.** Activity during UP versus DOWN phases of pupil self-regulation in the different regions of interest (ROIs). Statistical comparisons using paired-samples *t*-tests revealed significant effects (UP > DOWN) only in the LC ($t(21) = 3.84$; $p = 0.005$; $d = 0.82$; 95%CI$d$ [0.33;1.30]). Differences in the SN/VTA did not survive correction for multiple comparisons ($t(21) = 2.41$; $p = 0.10$; $d = 0.52$; 95%CI$d$ [0.06;0.96]). Differences in the NBM, SC and DRN were also not significant (all $p > 0.096$). **B.** Activity during UP versus DOWN phases of pupil self-regulation extracted from the control regions as depicted in Supplementary Figure 5A. Statistical comparisons using paired *t*-tests revealed no significant effects of pupil self-regulation (UP versus DOWN) on BOLD activity extracted from each of the masks at an uncorrected level (complete brainstem: $t(21) = 0.55$; $p = 0.59$; $d = 0.12$; 95%CI$d$ [-0.30;0.54]; 4th ventricle: $t(21) = 1.50$; $p = 0.15$; $d = 0.32$; 95%CI$d$ [-0.12;0.75]; all *p*-values uncorrected). **C.** To investigate post-hoc whether the significant effects observed in the LC during up- as compared to downregulation exceed those in the complete brainstem, we compared the difference scores (UP>rest minus DOWN>rest: UP-DOWN) between these regions. Paired-samples *t*-tests revealed that the difference in BOLD activity between UP and DOWN phases in the LC is indeed significantly larger than in the complete brainstem ($t(21) = 3.79$; $p = 0.001$; $d = 0.81$; 95%CI$d$ [0.32;1.29]). **D.** Correlations between continuous pupil size changes and BOLD response changes shown as z-values for the different ROIs. Statistical comparison (against 0) revealed significant effects for the SN/VTA ($t(21) = 4.54$; $p = 0.005$; $d = 0.97$; 95%CI$d$ [0.45;1.47]) and DRN ($t(21) = 3.22$; $p = 0.016$; $d = 0.69$; 95%CI$d$ [0.21;1.15]) . The NBM did not survive correction for multiple comparisons ($p = 0.051$) and for the LC (only trend-level; $p = 0.08$) and the SC ($p = 0.53$), statistics did not reach significance. All ROI analyses except for control analyses in B were corrected for multiple comparisons using sequential Bonferroni correction. Squares represent individual data. Error bars indicate SEM. Locus coeruleus (LC); substantia nigra (SN) and ventral tegmental area (VTA) combined into one ROI (SN/VTA); dorsal raphe nucleus (DRN); nucleus basalis of Meynert (NBM); and superior colliculus (SC).

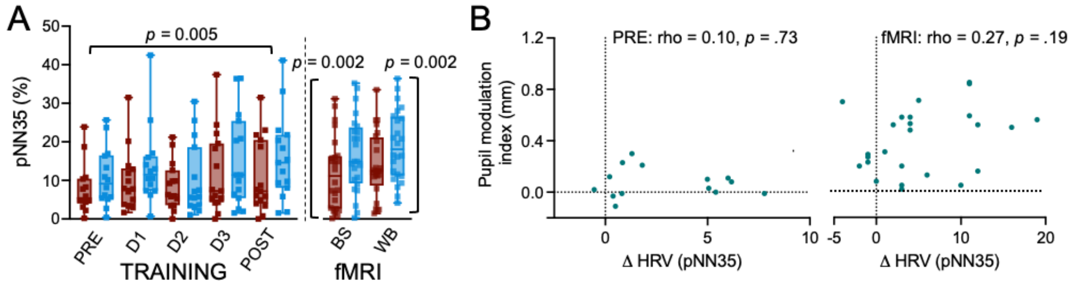

**Supplementary Figure 7.** Effects of pupil self-regulation on a different measure of heart rate variability (HRV). **A.** HRV was estimated as the percentage of successive R-R-intervals greater than 35ms (pNN35) and averaged for UP and DOWN trials across all participants for pupil-BF training (n = 14 for Pre no feedback and Day 1; n = 15 for all other sessions; left panel) and fMRI sessions (n = 24, right panel). Self-regulation of pupil size systematically modulated pNN35 with significantly larger values during DOWN than UP trials during pupil-BF training (significant difference between UP and DOWN trials when averaged across all sessions: Wilcoxon signed rank test: z = 2.84; *p* = 0.005; r = 0.76; but no significant learning effect when difference scores (DOWN-UP) for each session were tested using a robust repeated-measures ANOVA: $F(3.13, 28.21) = 1.18$; *p* = 0.33) and fMRI sessions (whole-brain Wilcoxon signed rank test: z = 3.06; *p* = 0.002; r = 0.62; brainstem Wilcoxon signed rank test: z = 3.37; *p* = 0.002; r = 0.69; sequential Bonferroni corrected). **B.** Non-significant Spearman rho correlation coefficients for pupil modulation indices (UP-DOWN) and pNN35 differences (DOWN-UP) prior to (left panel) and after pupil-BF training during fMRI (right panel). Please note that even though the pNN35 revealed significant differences between up- and downregulation of pupil size, we are still cautious to over-interpret this data calculated on time windows as short as 15s, since RMSSD is still preferred over the pNN-xx by most researchers. This may be not only because of the good validation of the RMSSD for ultra-short-term measurements but also because the pNN-xx introduces a rather arbitrary thresholding. Error bars indicate SEM. BS = brainstem fMRI; WB = whole-brain fMRI.

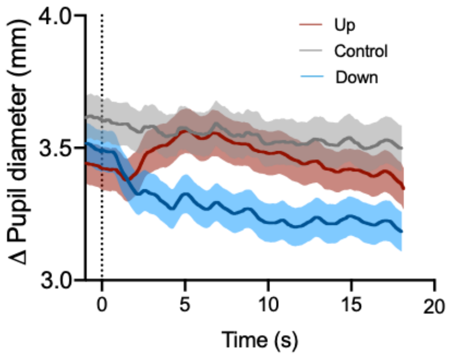

**Supplementary Figure 8.** Absolute pupil diameter time series during the pupil-BF oddball task averaged across participants for UP (red), DOWN (blue) and Control trials (grey) showing the last second of the baseline phase (-1-0s) as well as the complete modulation phase of 18s.

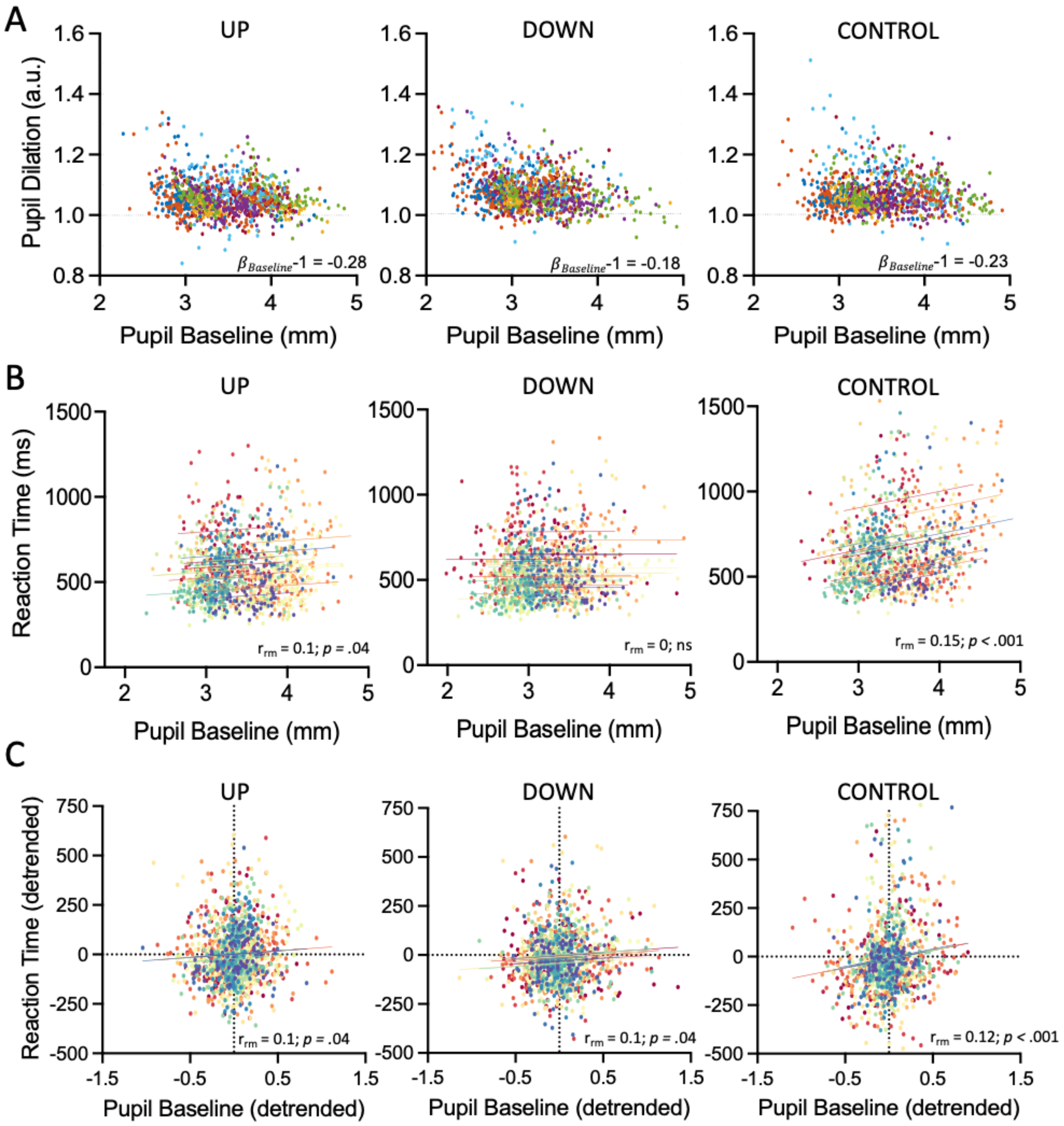

**Supplementary Figure 9.** Relationship of baseline pupil size with pupil dilation responses and reaction times to target tones during the auditory oddball task. **A.** Pupil dilation responses to target sounds and their relation to baseline pupil size. Since the correlation between a variable x and the variable y-x is biased, we conducted relative baseline correction of pupil dilation responses ($\frac{pupil\ dilation\ response\ peak}{baseline\ pupil\ size}$) and tested for an unbiased relationship by fitting the following model: $\beta_x$: $lm(log(y) \sim log(x))$.This model corresponds to the following equation: $log(y) = \beta_x log(x) + intercept + \epsilon$ which is equivalent to $\frac{y}{x} = x^{\beta x\text{-}1} \times e^{intercept} \times e^{\epsilon}$. Testing with a one-sample $t$-test for $\beta_x - 1 = 0$ (i.e., H0: baseline pupil size has no significant influence on pupil dilation responses), revealed indeed an inverse relationship between pupil dilation responses and their respective baseline pupil size averaged across 500ms before target onset during up- (left panel; $\beta_{\text{Baseline}}$ -1 = -0.28; $t(19)$ = -6.91, $p$ < .001, Cohen's $d$ = -1.55; 95%-CI$_d$ [-2.18; -0.88]), downregulation (middle panel; $\beta_{\text{Baseline}}$ -1 = -0.18; $t(19)$ = -6.74, $p$ < .001, Cohen's $d$ = -1.51; 95%-CI$_d$ [-2.14; -0.85]) and control trials (right panel; $\beta_{\text{Baseline}}$ -1 = -0.23; $t(19)$ = -4.85 $p$ < .001, Cohen's $d$ = -1.08; 95%-CI$_d$ [-1.63; -0.52])). **B.** Reaction times to target sounds were positively related to the respective baseline pupil size (averaged across 500ms before target onset) during upregulation (left panel) and control trials (right panel), but not during downregulation (middle panel) when taking the raw values measured during the task. **C.** When taking time-on-task effects into account by detrending the data (using the MATLAB function *detrend* with the method "linear" before submitting it to repeated measures correlations), significant relationships were revealed in all three conditions, even though the effect size of these effects were small showing only weak links between the measures. Similar analyses with pupil dilation responses instead of baseline pupil size did not reveal significant relationships to task performance indicated by reaction times (all $p > .10$).

**Supplementary Table 1.** Self-reported strategies during debriefing of all included participants used during the pupil-BF training. Please note that most participants in both groups imagined an emotional situation during the UP condition and a relaxing situation and focused on breathing during the DOWN condition.

| NF group | Up | Down |
|---|---|---|
| 1 | imagine getting injections | relaxing, changing breath, counting slowly |
| 2 | hate against opponent team (rivalry) | focussing on skiing/surfing |
| 3 | imaginge swearing/insulting/screaming at someone, imagine pain, positive/exciting memories | relaxing and letting go |
| 4 | running (Sprints) | Focus on breathing and imagining being at the beach hearing ocean sounds |
| 5 | Imagining happy moments, alternating with imagining me running | imagining sunlight |
| 6 | Imagining sports activity (e.g., running) | Imagining being in the sun |
| 7 | Imagining fearful situations (spiders and tight rooms) | Focused on breathing and how body feels while breathing (i.e., “scan” different body parts) |
| 8 | Imagining being nervous (e.g., important event) combined with sports activity (e.g., running) | “blank mind”, imagining situation just before falling asleep |
| 9 | Imagining stressful, very happy, aggressive moments | Focus on breathing and central fixation point, anything else is completely out of focus |
| 10 | Imagining fighting with someone and cursing in my head | Focused and concentrated breathing, let go of everything, just focus on myself |
| 11 | Imagining feeling of panic, stress, grief | Counting silently and imagining being tired lying in bed |
| 12 | Imagining emotional situations (spiders, verbally fighting with someone) combined with energization of behavior (running to not miss the train) | Think of nothing, “blank mind”, just focus on fixation point in the center |
| 13 | imagine anxious and painful situations, however I also felt like I may have breathed faster | positive thoughts, happy memories (visiting my family, for instance), relaxation (concentrating on breathing) |
| 14 | Imagine very positive and happy situations | Being mentally absent and focus solely on body |
| 15 | imagining sad situations | Using same strategies as when meditating, focusing on a particular point, concentration on breathing, having happy thoughts |
| 16 | Imagining moments with my family combined with physical activities (running/playing basketball) | focusing on my breath |
| 17 | imagined that the cabin is full of rats/spiders/insects, imagining that a clown is trying to kill me | “blank mind” |
| 18 | Imagining emotional situations (may have led to breathing more shortly) | not paying attention to the screen, focus on myself |
| 19 | imagining jumping from a cliff into the sea | imagined floating in the air, being weightless without purpose |
| 20 | Imagining scenes from a series | “blank mind”, think of nothing at all |
| 21 | Imagining stressful situation (exam), playing songs in my mind | Concentrate on fixation point in the center of the screen |
| 22 | Imagining spiders and running away from them | Breathing pattern, I use before falling asleep (4 - 7 - 8 (breathing in - stop – let go) |

| | | |
|---|---|---|
| 23 | Imagining stressful and disgusting situations (exams, failure, worms) | imagining favourite scenarios (holidays, food), focus on breathing |
| 24 | feeling stress: imagine an oral exam, insults or threats | “blank mind” |
| 25 | imagined doing pull-ups and tried to multiply large numbers in my head | imagined looking at the sun, focus on dot in center of the screen |
| 26 | Imagining being angry (shouting at someone), alternating with feeling of disgust and subtracting large numbers in my head | Think of nothing and focus on breathing |
| 27 | Imagining running to catch a bus, running away from someone | Focus on screen that is bright |
| 28 | Imaging happy situations (looking forward to a happy moment), imagining focussing in sports competition | Feel breathing, trying to relax and let go of everything |
| 29 | Imagine fight with sister alternating with running in sports | Focus on breathing, single body parts, imagine environment behind me |
| 30 | Imaging anxious and angry situations (being alone in the forest, shouting/cursing), singing in my head | Focus on breathing, everything is warm and bright in my body |
| 31 | Imagining insects on body | Think of nothing, just waves in my head, focus on body parts |
| 32 | Imagining very happy (surfing, snowboarding, reaching mountain top) and fearful moments (road accident) | Think of nothing |
| 33 | Imagining sad situations (death of relative) | Imagining falling asleep in bed |
| 34 | Imagining insects on body, feet, head | Focus on breathing, imagining lying in bed and relax |
| 35 | Imagining exciting situations (new research results) | Breasthing/clearing mind/ thinking its ok, increasing speed of word count during exercise |
| 36 | Imagining sports events (situations in soccer) and sad memories (break up), subtracting numbers | defocus (screen and environment) |
| 37 | Imagining emotional situations (pleasant like hugging and kissing and disgusting like worms and bugs) | Imagining yoga poses |
| 38 | Imagining pain when having an injury, combined with running | direktes Sonnenlicht, Strandatmosphäre, Fokus auf Finger |
| 39 | Imagining anxious situation (wasp sting, being allergic) | Imagining going to bed routine, being aware of body parts, especially seat bone on chair |
| 40 | mental weightlifting | focussing on breath, imagination of happy place, emptying the mind |
| 41 | Vividly imagining body in stress situation (mentally overreacting: “I need to get stressed”, "Go,go,go") | imagined myself sleeping in bed |
| 42 | Imagined spiders, snakes and bugs on my skin and running away from stranger with knife | Focused on breathing, imagined floating in water while not thinking of anything just focusing on body |
| 43 | Imagining tense and fearful situations | think of nothing |
| 44 | Thinking of fearful and disgusting situations (spiders, insects, running away from someone) | Imagining falling asleep |
| 45 | Imagining very heavy exercise | Focus on breathing and imagining relaxing situations |
| 46 | Imagining fearful situations (dogs) and running away | Imagining lying on balcony and watching/listening to mows, focus on own body positions |
| 47 | Imagining being chased in a haunted house | focusing on my breathing |
| 48 | Imagining very emotional situations (dancing and singing, giving birth) | Imagining green in forest, surfing in the ocean, meditating in piece and quiet |
| 49 | Imagining emotional situation (heavy fight in the past) | Imagining and feeling different body parts and being grounded with my feet while focusing on breathing |

| | | |
|---|---|---|
| 50 | thinking of very positive and happy situations | Imagining falling asleep and relaxed situations while focusing on body parts and breathing |
| 51 | Imagining joyful situations (pleasure during sports, being with friends and family) | Mentally counting (very slowly) |
| 52 | mentally running a route | Imagining going to a nice place and focus on breathing, alternating with thinking of nothing |
| **Control group** | **Up** | **Down** |
| 1 | Imagining stressful/sports situations | Focus on breathing, think of nothing |
| 2 | Imagining fighting with annoying person | Imagining relaxing music |
| 3 | shouting and arguing with a host-mate I don't like | Imagining lying under the fruit trees in my food forest at home |
| 4 | Imagining being angry, shouting at someone | Imagining that each body part is getting more and more heavy, watching my body from the outside; imagining lying in the sun and feeling it |
| 5 | Imagining sad, angry, happy, stressful (exam) situations | concentrated on how breathing happens |
| 6 | Imagining movie scenes | Imaging lying at the beach |
| 7 | Imagining very joyful moments | Focusing on breathing and point in the center of the screen |
| 8 | Imagining emotionally heavy situation (accident) and erotic situations | Imagining how I fall asleep, imagining noise in the background to think of nothing at all |
| 9 | Imagining spiders crawling on me, alternating with running | Imagining relaxing on beach and warm sun; imagining floating on water and falling asleep |
| 10 | Imagining hitting a punch bag | Focused on slow breathing |
| 11 | Imagining singing at a concert favorite song, kissing someone | thinking of the feeling before falling asleep (relaxed feeling) |
| 12 | Imagining situations where I was very angry, Imagining heavy exercise | Focus on breathing and imagining relaxing situations |
| 13 | Imagining fighting and shouting with someone | Day-dreaming |
| 14 | imagining stressful situations, panic and fear (may have led to more heavy breathing) | breathing slowly/deeply - imagining calm situations and landscapes - hugging someone |
| 15 | Imagining stressful situations with friends, imagined some high-action situations | Imagining being at the beach, watching sunset and shooting stars |
| 16 | Imagining stressful situations and heavy activity (more heavy breathing) | Focus on calm breathing, imagining relaxing and pleasant situation |
| 17 | Imagining fearful situations (dogs which are chasing me, someone pointing a rifle at me, chasing me) | Imagining being under water, getting a massage |
| 18 | I imagined the difficulties and situations that I passed through, something that life depends on | I imagined myself doing parasailing and relaxing travels with friends |
| 19 | Imagined pain in the toe, shoulder and back, and fearful/stressful situation (exam fear) | Focused on breathing and making it longer, repeating 'OM' in my mind |
| 20 | Thinking about the current Coronavirus; thinking about human rights situation in my; imagining spending time with my beautiful friend | Thinking of nothing, imagining myself lying in bed and looking at the sky |
| 21 | Imagining a very stressful situation (before an important speech, exams, failure) | Imagining a relaxing and calm place |

| | | |
|---|---|---|
| 22 | Imagining anxious situations (someone putting a spider on me, following me in the dark) , alternating with energization (exhausting end of a marathon) | Focus on breathing, imagine relaxing situation (getting a massage) |
| 23 | Imaging stressful and panic situations (exam, giving talk in front of many people, run away from murderer; painful injection) | Focus and slow down breathing; feeling heavy and solely focus on fingertips |
| 24 | Imagining emotional situations (whole range from love, fear, euphoria, hate/anger, panic, happiness) | Imagining relaxing situation or thinking of nothing at all |
| 25 | Imagining huge spider, being haunted or riding on a horse with full speed (mostly combination of everything) | Focusing on deep breathing, relaxing shoulders and listening to sounds in forest |
| 26 | Imagining emotional situation (i.e., heavy fight with someone, seeing a person I don't like) | Combining focus on breathing and trying to think of nothing with being aware of body (where do clothes touch the skin) |
| 27 | Imagining doing heavy exercises (run next to a steep slope, doing pushups) combined with emotional situation (stressful situations, having to go on stage soon, seeing a family member getting hurt) | Imagining lying in bed after tough training, imagining relaxing music in my head, getting a relaxing massage |

**Supplementary Table 2A.** Brainstem regions in addition to a-priori-defined regions of interest showing stronger activation during up- as compared to downregulation (UP > DOWN) in a mixed-effects analysis. Since there were no significant regions in the brainstem when implementing cluster-correction ($z > 2.3$; $p < 0.05$), all reported results refer to analyses at an uncorrected $p < .05$. We implemented a minimum cluster size of 10 voxels derived from the smallest mask of a recently published brainstem atlas covering all important brainstem nuclei involved in arousal and motor control (Bianciardi, 2021). Coordinates reported are peak coordinates in MNI space overlapping with anatomical labels determined using the Brainstem Navigator (Bianciardi, 2021).

| | **peak coordinate overlapping with brainstem region** | | | |
|---|---|---|---|---|
| **Region** | **x** | **y** | **z** | **z-value** |
| Inferior/<br>Superior Medullary Reticular Formation | 6<br>6 | -42<br>-44 | -58<br>-48 | 2.71<br>2.30 |
| Laterodorsal Tegmental Nucleus – Central Gray of the Rhomboencephalon | 2 | -38 | -28 | 3.00 |
| Microcellular Tegmental Nucleus | -8 | -32 | -10 | 2.91 |
| Periaqueductal gray | 0 | -30 | -4 | 2.37 |
| Viscero-sensory-motor Nuclei Complex | -2 | -42 | -48 | 2.57 |
| Pedunculopontine Nucleus | -8 | -28 | -12 | 2.61 |

**Supplementary Table 2B.** Brainstem regions in addition to mentioned regions of interest covarying with pupil diameter during pupil self-regulation in a mixed-effects analysis (cluster-corrected at $z > 2.3$; $p < .05$) of the brainstem fMRI session. Coordinates reported are peak coordinates in MNI space overlapping with anatomical labels determined using the Brainstem Navigator[1].

| | **peak coordinate overlapping with brainstem region** | | | |
|---|---|---|---|---|
| **Region** | **x** | **y** | **z** | **z-value** |

| | | | | |
|---|---|---|---|---|
| Inferior/<br>Superior Medullary Reticular Formation | -4<br>4 | -44<br>-40 | -52<br>-46 | 3.23<br>2.84 |
| Laterodorsal Tegmental Nucleus –<br>Central Gray of the Rhomboencephalon | 0 | -38 | -34 | 3.12 |
| Microcellular Tegmental Nucleus | 8 | -30 | -10 | 2.48 |
| Periaqueductal gray | 2 | -30 | -12 | 3.04 |
| Viscero-sensory-motor Nuclei Complex | -2 | -44 | -54 | 3.57 |
| Pedunculopontine Nucleus | 10 | -30 | -14 | 4.22 |
| Mesencephalic Reticular Formation | -4 | -26 | -4 | 2.89 |
| Parvicellular Reticular Nucleus | 8 | -38 | -44 | 3.20 |
| Paramedian Nucleus | 0 | -30 | -22 | 3.12 |
| Raphe Obscurus | 0 | -42 | -56 | 2.97 |
| Pontine Reticular Nucleus | 0 | -30 | -22 | 3.12 |

**Supplementary Table 3A.** Activation clusters, corresponding size, anatomical region, FWE-corrected *p*-value, peak coordinate in MNI space, and maximum z-value for the up > down contrast thresholded at z > 3.1 and FWE-corrected using a cluster significance level of $p < .05$ from the whole-brain fMRI session. Reported anatomical labels were determined using the Jülich Histological[2], the Harvard-Oxford cortical and subcortical structural[3,4] and probabilistic cerebellar atlases[5], and correspond to the location of maxima within each cluster.

| | | | | **peak coordinate** | | | |
|---|---|---|---|---|---|---|---|
| **cluster** | **# voxels** | **Region of peak** | **pFWEcorr.** | **x** | **y** | **z** | **z-value** |
| 1 | 5916 | Paracingulate/Anterior Cingulate Cortex | <.001 | -2 | 20 | 38 | 5.39 |
| 2 | 5129 | Cerebellum (VI; Crus I) | <.001 | 32 | -58 | -28 | 5.73 |
| 3 | 1111 | Globus Pallidus | <.001 | -16 | -6 | -4 | 5.11 |
| 4 | 656 | Praecuneus | <.001 | -8 | -56 | 58 | 4.51 |
| 5 | 570 | Primary motor/somatosensory cortex | <.001 | 30 | -28 | 58 | 5.09 |

| | | | | | | | |
|---|---|---|---|---|---|---|---|
| 6 | 520 | Dorsolateral Prefrontal Cortex | <.001 | -26 | 44 | 26 | 4.63 |
| 7 | 269 | Caudate Nucleus | <.001 | 20 | 8 | 18 | 4.89 |

**Supplementary Table 3B.** Activation clusters, corresponding size, anatomical region, FWE-corrected *p*-value, peak coordinate in MNI space, and maximum z-value for the down > up contrast thresholded at z > 3.1 and FWE-corrected using a cluster significance level of $p < .05$ from the whole-brain fMRI session. Reported anatomical labels were determined using the Jülich Histological Atlas[2] and correspond to the location of maxima within each cluster.

| | | | | **peak coordinate** | | | |
|---|---|---|---|---|---|---|---|
| **cluster** | **# voxels** | **Region of peak** | **pFWEcorr.** | **x** | **y** | **z** | **z-value** |
| 1 | 716 | Visual Cortex V5 | <.001 | -46 | -82 | 4 | 4.42 |

**Supplementary Table 4.** Activation clusters, corresponding size, anatomical region, FWE-corrected *p*-value, peak coordinate in MNI space, and maximum z-value covarying with pupil size thresholded at z > 3.1 and FWE-corrected using a cluster significance level of $p < .05$ from the whole-brain fMRI session. Please note that pupil time series were shifted by 1s to account for delays between brain and pupillary signals. Reported anatomical labels were determined using the Harvard-Oxford cortical and subcortical structural[3,4], the probabilistic cerebellar atlases[5], and the Talairach atlas registered to MNI152 standard space[6], and correspond to the location of maxima within each cluster.

| | | | | **peak coordinate** | | | |
|---|---|---|---|---|---|---|---|
| **cluster** | **# voxels** | **Region of peak** | **pFWEcorr.** | **x** | **y** | **z** | **z score** |
| 1 | 2163 | Midbrain | <.001 | 4 | -16 | -14 | 5.50 |
| 2 | 1134 | Anterior Cingulate Cortex | <.001 | -2 | 30 | 26 | 5.50 |
| 3 | 1014 | Cerebellum (VI, Crus I) | <.001 | 32 | -58 | -36 | 4.87 |
| 4 | 184 | Cerebellum (Vermis) | 0.02 | -2 | -14 | 14 | 4.79 |
| 5 | 148 | Precuneus | 0.04 | -8 | -54 | 58 | 3.93 |

**Supplementary Table 5.** Description of group means and standard deviation for data presented in the main manuscript.

| **Figure** | **Group** | **Measure** | **Mean** | **SD** |
|---|---|---|---|---|
| 2C | Pupil-BF | Pupil modulation index Day 1<br>Pupil modulation index Day 2<br>Pupil modulation index Day 3<br>Pupil modulation index post no feedback | 0.20<br>0.26<br>0.29<br>0.34 | 0.17<br>0.20<br>0.25<br>0.28 |
| 2C | Control I | Pupil modulation index Day 1<br>Pupil modulation index Day 2<br>Pupil modulation index Day 3<br>Pupil modulation index post no feedback | 0.04<br>0.04<br>0.08<br>0.09 | 0.08<br>0.14<br>0.14<br>0.12 |
| SF 3C | Control II | Pupil modulation index Day 1<br>Pupil modulation index Day 2<br>Pupil modulation index Day 3<br>Pupil modulation index Day 4 | 0.06<br>0.06<br>0.06<br>0.09 | 0.09<br>0.10<br>0.11<br>0.10 |

| | | | | |
|---|---|---|---|---|
| SF 1A | Pupil-BF | Pupil Modulation Up Day 1 | 0.14 | 0.11 |
| | | Pupil Modulation Up Day 2 | 0.18 | 0.11 |
| | | Pupil Modulation Up Day 3 | 0.15 | 0.13 |
| | | Pupil Modulation Up no feedback post | 0.10 | 0.15 |
| | | Pupil Baseline Up Day 1 | 0.00 | 0.00 |
| | | Pupil Baseline Up Day 2 | 0.00 | 0.00 |
| | | Pupil Baseline Up Day 3 | 0.00 | 0.00 |
| | | Pupil Baseline Up no feedback post | 0.00 | 0.00 |
| SF 1A | Control | Pupil Modulation Up Day 1 | -0.00 | 0.10 |
| | | Pupil Modulation Up Day 2 | 0.02 | 0.14 |
| | | Pupil Modulation Up Day 3 | 0.04 | 0.12 |
| | | Pupil Modulation Up no feedback post | 0.01 | 0.12 |
| | | Pupil Baseline Up Day 1 | 0.00 | 0.00 |
| | | Pupil Baseline Up Day 2 | 0.00 | 0.00 |
| | | Pupil Baseline Up Day 3 | 0.00 | 0.00 |
| | | Pupil Baseline Up no feedback post | 0.00 | 0.00 |
| SF 1B | Pupil-BF | Pupil Modulation Down Day 1 | -0.06 | 0.18 |
| | | Pupil Modulation Down Day 2 | -0.08 | 0.15 |
| | | Pupil Modulation Down Day 3 | -0.14 | 0.18 |
| | | Pupil Modulation Down no feedback post | -0.24 | 0.18 |
| | | Pupil Baseline Down Day 1 | 0.00 | 0.00 |
| | | Pupil Baseline Down Day 2 | 0.00 | 0.00 |
| | | Pupil Baseline Down Day 3 | 0.00 | 0.00 |
| | | Pupil Baseline Down no feedback post | 0.00 | 0.00 |
| SF 1B | Control | Pupil Modulation Down Day 1 | -0.04 | 0.09 |
| | | Pupil Modulation Down Day 2 | -0.02 | 0.12 |
| | | Pupil Modulation Down Day 3 | -0.04 | 0.13 |
| | | Pupil Modulation Down no feedback post | -0.07 | 0.09 |
| | | Pupil Baseline Down Day 1 | 0.00 | 0.00 |
| | | Pupil Baseline Down Day 2 | 0.00 | 0.00 |
| | | Pupil Baseline Down Day 3 | 0.00 | 0.00 |
| | | Pupil Baseline Down no feedback post | 0.00 | 0.00 |
| SF 2A | Pupil-BF | Pupil Baseline Up Day 1 | 3.58 | 0.55 |
| | | Pupil Baseline Up Day 2 | 3.53 | 0.53 |
| | | Pupil Baseline Up Day 3 | 3.56 | 0.61 |
| | | Pupil Baseline Up no feedback post | 3.50 | 0.58 |
| | | Pupil Baseline Down Day 1 | 3.57 | 0.55 |
| | | Pupil Baseline Down Day 2 | 3.52 | 0.53 |
| | | Pupil Baseline Down Day 3 | 3.55 | 0.61 |
| | | Pupil Baseline Down no feedback post | 3.55 | 0.56 |
| SF 2A | Control I | Pupil Baseline Up Day 1 | 3.53 | 0.54 |
| | | Pupil Baseline Up Day 2 | 3.52 | 0.53 |
| | | Pupil Baseline Up Day 3 | 3.51 | 0.43 |
| | | Pupil Baseline Up no feedback post | 3.49 | 0.44 |
| | | Pupil Baseline Down Day 1 | 3.54 | 0.56 |
| | | Pupil Baseline Down Day 2 | 3.52 | 0.57 |
| | | Pupil Baseline Down Day 3 | 3.51 | 0.44 |
| | | Pupil Baseline Down no feedback post | 3.50 | 0.50 |
| SF 2B | Control II | Pupil Baseline Up Day 1 | 3.49 | 0.50 |
| | | Pupil Baseline Up Day 2 | 3.49 | 0.38 |
| | | Pupil Baseline Up Day 3 | 3.52 | 0.41 |
| | | Pupil Baseline Up no feedback post | 3.47 | 0.33 |
| | | Pupil Baseline Down Day 1 | 3.47 | 0.51 |
| | | Pupil Baseline Down Day 2 | 3.46 | 0.43 |
| | | Pupil Baseline Down Day 3 | 3.50 | 0.41 |
| | | Pupil Baseline Down no feedback post | 3.50 | 0.37 |
| SF 2C | Pupil-BF | Pupil Baseline Up Brainstem | 3.63 | 0.54 |
| | | Pupil Baseline Down Brainstem | 3.65 | 0.56 |
| | | Pupil Baseline Up Whole-Brain | 3.65 | 0.54 |

| | | | | |
|---|---|---|---|---|
| | | Pupil Baseline Down Whole-Brain | 3.70 | 0.56 |
| SF 2D | Pupil-BF | Pupil Baseline Up Oddball<br>Pupil Baseline Down Oddball<br>Pupil Baseline Control Oddball | 3.44<br>3.50<br>3.62 | 0.36<br>0.35<br>0.38 |
| 3B | Pupil-BF | z-value LC up<br>z-value LC down<br>z-value SN/VTA up<br>z-value SN/VTA down<br>z-value SC up<br>z-value SC down<br>z-value DRN up<br>z-value DRN down<br>z-value NBM up<br>z-value NBM down | 0.15<br>-0.10<br>0.07<br>-0.09<br>-0.23<br>-0.45<br>-0.01<br>-0.08<br>0.43<br>0.03 | 0.51<br>0.54<br>0.32<br>0.26<br>0.56<br>0.49<br>0.40<br>0.42<br>0.75<br>0.67 |
| 3C | Pupil-BF | z-value LC<br>z-value SN/VTA<br>z-value NBM<br>z-value DRN<br>z-value SC | 0.31<br>0.44<br>0.50<br>0.31<br>0.17 | 0.56<br>0.41<br>0.73<br>0.54<br>0.73 |
| SF 5B | Pupil-BF | z-value complete brainstem up<br>z-value complete brainstem down<br>z-value 4th ventricle up<br>z-value 4th ventricle down | 0.00<br>-0.08<br>-0.02<br>-0.03 | 0.17<br>0.16<br>0.60<br>0.49 |
| SF 5C | Pupil-BF | z-value LC up-down<br>z-value SN/VTA up-down<br>z-value NBM up-down<br>z-value complete brainstem up-down | 0.25<br>0.16<br>0.40<br>0.08 | 0.34<br>0.26<br>0.70<br>0.20 |
| SF 6A | Pupil-BF | z-value LC up<br>z-value LC down<br>z-value SN/VTA up<br>z-value SN/VTA down<br>z-value SC up<br>z-value SC down<br>z-value DRN up<br>z-value DRN down<br>z-value NBM up<br>z-value NBM down | 0.09<br>-0.06<br>0.03<br>-0.02<br>-0.07<br>-0.11<br>-0.01<br>-0.06<br>0.19<br>0.07 | 0.26<br>0.27<br>0.10<br>0.10<br>0.15<br>0.23<br>0.18<br>0.19<br>0.32<br>0.27 |
| SF 6B | Pupil-BF | z-value complete brainstem up<br>z-value complete brainstem down<br>z-value 4th ventricle up<br>z-value 4th ventricle down | -0.00<br>-0.01<br>0.02<br>-0.06 | 0.07<br>0.05<br>0.22<br>0.22 |
| SF 6C | Pupil-BF | z-value LC up-down<br>z-value complete brainstem up-down | 0.15<br>0.01 | 0.19<br>0.07 |
| SF 6D | Pupil-BF | z-value LC<br>z-value SN/VTA<br>z-value NBM<br>z-value DRN<br>z-value SC | 0.12<br>0.13<br>0.24<br>0.16<br>0.03 | 0.31<br>0.14<br>0.43<br>0.23<br>0.24 |
| 5A | Pupil-BF | Heart Rate pre training no feedback up<br>Heart Rate Day 1 up<br>Heart Rate Day 2 up<br>Heart Rate Day 3 up<br>Heart Rate post training no feedback up<br>Heart Rate pre training no feedback down<br>Heart Rate Day 1 down<br>Heart Rate Day 2 down<br>Heart Rate Day 3 down | 84.22<br>82.02<br>83.21<br>80.69<br>79.98<br>82.04<br>78.35<br>79.27<br>75.79 | 12.14<br>10.52<br>11.05<br>12.17<br>12.60<br>11.30<br>10.46<br>10.08<br>11.06 |

| | | | | |
|---|---|---|---|---|
| | | Heart Rate post training no feedback down | 74.54 | 10.65 |
| 5A | Pupil-BF | Heart Rate brainstem fMRI up<br>Heart Rate brainstem fMRI down<br>Heart Rate whole-brain fMRI up<br>Heart Rate whole-brain fMRI down | 79.07<br>73.60<br>76.92<br>70.18 | 13.52<br>11.91<br>7.67<br>8.03 |
| 5D | Pupil-BF | RMSSD pre training no feedback down-up<br>RMSSD Day 1 down-up<br>RMSSD Day 2 down-up<br>RMSSD Day 3 down-up<br>RMSSD post training no feedback down-up | 1.10<br>-2.77<br>0.00<br>3.08<br>1.71 | 4.82<br>13.42<br>7.72<br>15.18<br>16.52 |
| 5D | Pupil-BF | RMSSD brainstem fMRI down-up<br>RMSSD whole-brain fMRI down-up | 5.63<br>6.79 | 17.16<br>19.60 |
| SF 3D | Control II | Heart Rate Day 1 up<br>Heart Rate Day 1 down<br>Heart Rate Day 2 up<br>Heart Rate Day 2 down<br>Heart Rate Day 3 up<br>Heart Rate Day 3 down<br>Heart Rate post training no feedback up<br>Heart Rate post training no feedback down | 78.20<br>76.13<br>80.12<br>78.66<br>78.43<br>76.52<br>75.22<br>73.19 | 11.65<br>11.27<br>9.14<br>9.13<br>7.50<br>7.68<br>8.20<br>9.04 |
| SF 3D | Control II | RMSSD Day 1 up<br>RMSSD Day 1 down<br>RMSSD Day 2 up<br>RMSSD Day 2 down<br>RMSSD Day 3 up<br>RMSSD Day 3 down<br>RMSSD post training no feedback up<br>RMSSD post training no feedback down | 40.55<br>40.84<br>46.06<br>44.64<br>47.95<br>49.74<br>39.30<br>41.11 | 25.91<br>26.91<br>48.84<br>49.17<br>52.32<br>53.47<br>24.83<br>24.47 |
| 6C | Pupil-BF | Baseline-corrected pupil diameter up<br>Baseline-corrected pupil diameter down<br>Baseline-corrected pupil diameter control | 0.03<br>-0.23<br>-0.07 | 0.13<br>0.17<br>0.08 |
| 6E | Pupil-BF | Reaction time to targets up<br>Reaction time to targets down<br>Reaction time to targets control | 578.40<br>551.72<br>639.20 | 104.81<br>96.68<br>125.60 |
| 6E | Pupil-BF | SD of reaction time to targets up<br>SD of reaction time to targets down<br>SD of reaction time to targets control | 0.18<br>0.17<br>0.20 | 0.07<br>0.07<br>0.07 |
| SF 7A | Pupil-BF | pNN35 pre training no feedback up<br>pNN35 pre training no feedback down<br>pNN35 Day 1 up<br>pNN35 Day 1 down<br>pNN35 Day 2 up<br>pNN35 Day 2 down<br>pNN35 Day 3 up<br>pNN35 Day 3 down<br>pNN35 post training no feedback up<br>pNN35 post training no feedback down | 7.86<br>10.77<br>9.97<br>13.72<br>8.58<br>10.14<br>11.81<br>14.98<br>10.95<br>15.73 | 6.47<br>7.61<br>8.02<br>10.15<br>6.12<br>9.33<br>10.51<br>11.99<br>9.57<br>11.04 |
| SF 7A | Pupil-BF | pNN35 brainstem fMRI up<br>pNN35 brainstem fMRI down<br>pNN35 whole-brain fMRI up<br>pNN35 whole-brain fMRI down | 11.80<br>16.84<br>13.88<br>19.06 | 9.35<br>9.79<br>8.55<br>9.08 |
